# GATA2 SUMOylation Regulates GATA2 Stability to Control Uterine Serous Carcinoma Invasion

**DOI:** 10.64898/2026.11.12.765417

**Authors:** Anuoluwapo A. Mattix, Paula M. Mañán-Mejías, Usha Polaki, Apoorva T. Patil, Finnbar Reed-McBain, Peng Liu, Molly A. Accola, William M. Rehrauer, María Virumbrales-Muñoz, Daniel R Matson

## Abstract

Uterine serous carcinoma (USC) is an aggressive uterine cancer subtype that rapidly invades the uterine myometrium and is responsible for 40% of uterine cancer deaths. The molecular mechanisms that underlie this aggressive phenotype are poorly understood. We reported recently that USC tumors with low expression of the zinc-finger transcription factor GATA Binding Protein 2 (GATA2) exhibit aggressive tissue invasion, while retention of GATA2 expression is an independent predictor of 100% recurrence-free and cancer-specific survival in patients with early-stage USC. Here, we explored the mechanism of GATA2 loss in USC tumors that portend invasion. We show that GATA2 in human USC cell lines has a rapid 128-minute protein half-life due to post-translational SUMOylation, which targets GATA2 for ubiquitin-dependent proteasomal degradation. Treatment with small-molecule SUMOylation inhibitors, siRNA-mediated depletion of SUMO-2/3, or mutation of GATA2 K222 and K389 increases GATA2 levels and suppresses invasion in USC cell lines and USC tumor spheroids without impacting cell proliferation or viability. Primary USC tumors from newly diagnosed patients show markedly increased SUMO-2/3 levels and upregulation of SUMOylation pathways compared to other uterine cancer subtypes or benign endometrium, and heat-shock mediated induction of USC SUMO signaling leads to a SUMOylation-dependent reduction in GATA2 levels. Our results implicate SUMOylation as a dynamic regulator of USC GATA2 levels and support a model in which stress-related increases in SUMO activity deplete GATA2 to drive USC invasion. Our findings also raise the possibility that small-molecule SUMOylation inhibitors may benefit patients with early-stage USC disease.

## Introduction

Uterine serous carcinoma (USC) is a highly aggressive subtype of uterine cancer that accounts for only 10% of total uterine cancer cases but is responsible for 40% of uterine cancer deaths. The incidence of USC has been increasing by 4.9% annually for several decades, which largely accounts for the rise in the number of total uterine cancer diagnoses, estimated to reach 68,270 new U.S. cases in 2026.^1,2^ The survival statistics for USC have not improved over this same period, and the rising incidence and poor treatment response of USC now drive the surge in uterine cancer mortality, estimated to reach 14,450 uterine cancer deaths in the U.S. in 2026.^2,3^

Several factors contribute to high USC mortality, but chief among them is the predilection for USC to rapidly invade into the uterine myometrium and metastasize to other tissues. Half of USC patients have metastatic (extra-uterine) disease at diagnosis, defined by the International Federation of Gynecology and Obstetrics (FIGO) as stages II-IV, and experience poor outcomes.^4^ Even among USC patients with tumors apparently localized to the uterus at diagnosis (FIGO Stage I), 30% will experience disease recurrence after staging hysterectomy and have a median survival of 10-25 months, with nearly all dying of their cancer.^5–10^ USC recurrence is attributed to small numbers of invasive tumor cells that have escaped from the uterus but evaded detection at diagnosis, highlighting the rapid and unpredictable pattern of USC invasion and spread.^5^ Unfortunately, the upstream molecular pathways that promote USC invasion are poorly understood and therefore with the exception of Herceptin which benefits a subset of USC patients we lack targeted interventions to suppress tumor growth and spread.

The pathogenic events that underlie USC are slowly emerging. USC is genomically defined by early and near-universal *TP53* inactivation, followed by acquisition of high levels of aneuploidy, and less frequently, *ERBB2* amplification and/or mutations in *PIK3CA*, *PPP2R1A*, and *FBXW7*, and *CCNE1*.^11^ However, these abnormalities have been principally mapped to cell-cycle, PI3K, and MAPK signaling rather than pro-invasion pathways.^12^ *HER2/ERBB2* overexpression or amplification also occurs in 25% to 30% of USC tumors with downstream PI3K/AKT and MAPK signaling. In these patients with HER2^+^ tumors, anti-HER2 agents (trastuzumab) can be administered to suppress invasion and promote progression-free survival.^13^ USCs retain the pro-invasion adhesion molecule E-cadherin (38% of tumors) far less compared to endometrioid carcinomas (95% of tumors), and instead L1 cell adhesion molecule (L1CAM) is expressed in ∼77% of USCs with its expression tracking lymphovascular invasion and distant recurrence.^14,15^ In uterine cancer cell lines, TGF-β1 can induce L1CAM and downregulate E-cadherin in a Slug (SNAI2)-dependent manner.^16^ Critically, although the overexpression of HER2 and L1CAM can drive USC invasion, these events fail to explain why USC tumors of the same size with the same apparent cytogenetic background and HER2 and L1CAM expression profiles may be non-invasive or highly metastatic at the time of diagnosis.

GATA Binding Protein 2 (GATA2) is a zinc finger transcription factor that is expressed in the healthy uterus.^17,18^ The progesterone receptor (PGR) binds the *GATA2* promoter to drive *GATA2* transcription, while GATA2 also binds the *PGR* promoter to drive *PGR* transcription, forming a GATA2-PGR auto-regulatory positive feedback loop.^19^ GATA2 then co-localizes with PGR at PGR-target genes to facilitate the expression of PGR-driven transcriptional programs. In doing so, GATA2 critically enables important uterine functions including implantation of the fertilized embryo. We recently demonstrated that depletion of GATA2 drives USC invasion *in vitro*.^20^ Reduced GATA2 expression in clinical samples of primary USC tumors also closely correlates with myometrial invasion, lymphovascular invasion, and metastasis.^20^ Across a multi-institutional retrospective cohort of n=127 low-stage USC patients, we used routine GATA2 immunohistochemistry (IHC) to identify 33% of patients with tumors retaining GATA2 expression. These patients had 100% recurrence-free survival and 100% disease-specific survival over a median follow-up period of 3.2 years. In contrast, recurrence-free survival was only 60% in patients with USC tumors deficient in GATA2. Collectively, GATA2 expression was a positive and independent predictor of cancer-related survival, even in patients who received no adjuvant chemotherapy. Conversely, the loss of GATA2 predicted USC invasion regardless of tumor size, chemotherapy response, HER2 expression, *TP53* status, level of chromosome instability, and other metrics.^20^

Here, we designed studies to elucidate the molecular mechanisms that dictate GATA2 levels in USC and thereby determine tumor invasion and patient outcome. We show that USC GATA2 levels are subject to both transcriptional and post-translational regulation, but post-translational regulation dominates in patient tumors. The rapid turnover of GATA2 in cultured human USC cells is driven by SUMOylation, which targets the protein for ubiquitin-dependent proteasome-mediated degradation. Small-molecule inhibition of SUMOylation or expression of GATA2 SUMOylation mutants prolongs GATA2 half-life, increases GATA2 levels, and suppresses USC invasion independently of cell proliferation or survival in both USC cell lines and USC tumor spheroids. Primary USC tumors from treatment-naïve patients have elevated SUMO protein levels and enhanced SUMOylation pathway activity compared to either benign endometrium or uterine endometrioid carcinoma (UEC). Finally, stress-induced induction of SUMOylation pathways depletes GATA2 in USC cell lines by a process that can be blocked with small-molecule SUMOylation inhibitors. Our findings support a paradigm in which induction of stress-related SUMOylation pathways triggers the degradation of GATA2 and enables USC invasion, and implicate GATA2 stabilization as a candidate therapeutic approach for USC.

## Results

### Post-translational mechanisms predominantly determine GATA2 levels in USC tumors

PGR directly promotes *GATA2* transcription in healthy endometrium.^19^ To test whether the expression of GATA2 protein in USC is primarily determined by transcriptional regulation, we evaluated GATA2 protein levels by western blot and *GATA2* mRNA levels by RT-PCR across 7 patient-derived USC cell lines (Fig 1A-B). This revealed a significant positive, linear relationship between GATA2 protein and mRNA levels. We reported earlier that 33% of low stage (FIGO stage I) USC tumors have retained GATA2 protein expression (GATA2^high^) while 67% have reduced or absent expression (GATA2^low^), and low GATA2 strongly predicts USC myometrial invasion and worse survival. We reasoned that if *GATA2* mRNA levels correlate with GATA2 protein levels in patient USC tumors, they also should strongly predict USC recurrence and tumor invasion. We isolated all n=24 low stage (FIGO stage I) USCs from The Cancer Genome Atlas (TCGA) and classified the 8 tumors with highest *GATA2* RNA expression as RNA High, and the remaining 16 tumors as GATA2 Low (Fig 1C). The uterine cancer TCGA dataset lacks a linked quantitative proteomics dataset and so comparisons to GATA2 protein levels were not possible. However, TCGA has the most USC cases and most complete clinical outcome data of publicly available datasets, rendering it optimal for our study. TCGA USC *GATA2* RNA levels showed a strong negative correlation with *GATA2* gene methylation (Fig 1D), suggesting that methylation may be a central mechanism regulating *GATA2* USC transcription. However, unlike GATA2 protein, *GATA2* RNA levels did not predict USC recurrence and did not correlate with tumor invasion (Fig 1E-F).

**Figure 1:**
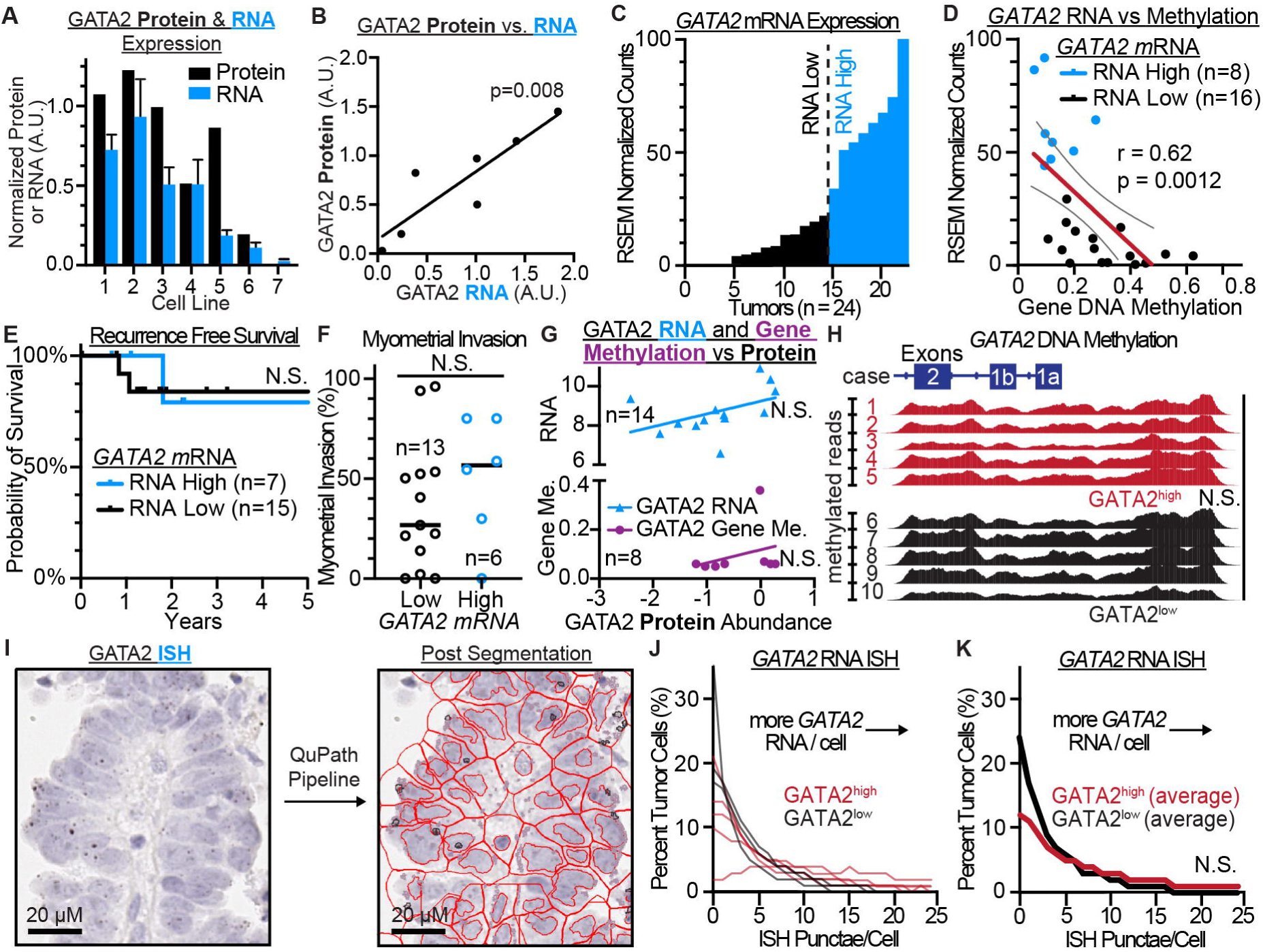
GATA2 proteins levels are subject to both transcriptional and post-translational regulation, but post-translational mechanisms dominate in patient tumors. **(A-B)** GATA2 protein levels plotted against *GATA2* mRNA levels in 7 patient-derived USC cell lines. **(C)** Histogram of *GATA2* mRNA levels across stage I USC tumors in TCGA. Blue bars indicate top 33% of cases based on *GATA2* mRNA level (RNA High). **(D)** *GATA2* RNA levels plotted against *GATA2* gene DNA methylation in TCGA. Blue circles indicate top 33% of cases based on *GATA2* mRNA level (RNA High). **(E)** Recurrence free survival and **(F)** myometrial invasion of stage I GATA2 RNA High and RNA Low cases from TCGA. **(G)** *GATA2* mRNA and gene methylation plotted against GATA2 protein levels derived from USCs in the CPTAC database. **(H)** Methylation across the *GATA2* proximal promoter through exon 2 in tumors scored as either GATA2^high^ or GATA2^low^ by GATA2 immunohistochemistry. **(I-K)** GATA2 in situ hybridization quantiying *GATA2* mRNA particles per cell in GATA2^high^ and GATA2^low^ USCs. **(I)** Example of GATA2 ISH slide and cell segmentation algorithm for quantifying ISH puncta per cell. **(J-K)** *GATA2* ISH puncta per cell plotted as **(J)** individual cases and **(K)** average across GATA2^high^ and GATA2^low^ cases.

These results suggested that while GATA2 RNA and protein closely correlate in USC cell lines, this relationship does not extend to patient USC tumors. We confirmed this finding by isolating multi-omic USC datasets from the Clinical Proteomic Tumor Analysis Consortium (CPTAC) database.^21^ While the uterine cancer discovery CPTAC dataset includes only n=14 USC tumors, these include tumor-matched gene methylation, transcriptomics, and quantitative proteomics data. The CPTAC dataset also includes benign endometrium controls. We first correlated GATA2 protein levels to both *GATA2* mRNA and *GATA2* gene methylation, and in both cases found no significant correlation (Fig 1G). Next, we evaluated the relationship between PGR and GATA2, which are known to co-regulate each other’s transcription in benign endometrium.^19^ We previously reported that PGR and GATA2 protein levels correlated in USC tumors within a multi-institutional USC tumor cohort,^20^ similarly found a highly significant positive correlation in both benign endometrium (p=0.0004, Supplementary Fig 1A) and USC tumors (p=0.0025, Supplementary Fig 1B) within CPTAC cases. We therefore hypothesized that PGR was continuing to regulate *GATA2* transcription in USC. However, USC PGR protein levels showed no statistical correlation with *GATA2* mRNA levels, as would be expected if PGR was acting as a transcriptional regulator of *GATA2* (p=0.46, Suppl Fig 1C). In contrast, USC GATA2 protein levels positively correlated with *PGR* mRNA levels (p=0.015, Supplementary Fig 1D). Therefore, analyses from publicly available datasets strongly argue that GATA2 protein levels are not predominantly determined by *GATA2* mRNA levels. They also suggest that PGR is not a primary regulator of USC *GATA2* transcription. Loss of PGR expression is characteristic of USC,^22^ and while candidate mechanisms including promoter hypermethylation, PRC2-mediated silencing, and miR-96 repression have been described in endometrial cancer cell lines,^23^ none has been validated in primary USC, and the upstream event driving PGR silencing remains unidentified.

The CPTAC database measures GATA2 levels by quantitative mass spectrometry, while we evaluated GATA2 levels in patient tumors across multiple institutions using IHC.^20^ Therefore, we sought to further confirm that GATA2 RNA and protein levels did not correlate in USC when assessed by IHC in our large multi-institutional patient cohort. Initially, because *GATA2* CpG methylation is known to silence *GATA2* expression in endometriosis,^24^ gastric cancer,^25^ and acute myeloid leukemia,^26^ we selected 5 GATA2^low^ USCs and 5 GATA2^high^ USCs from our published patient cohort and mapped DNA methylation across the *GATA2* promoter and through exon 2 (Fig 1H). This revealed no difference in *GATA2* methylation levels between GATA2^low^ and GATA2^high^ USCs. Next, because GATA2 protein levels are scored as percent positive tumor cell nuclei, we confirmed we were not missing positive RNA-protein correlations in single tumor cells because of bulk RNA sequencing approaches. These studies subjected the same tissue blocks from the 10 tumors used in our GATA2 IHC and *GATA2* gene methylation analysis to *GATA2 in situ* hybridization and uniformly applied a computational pipeline across the tumor regions to score the number of RNA ISH punctae in thousands of individual cells using QuPath (Fig 1I).^27^ When we plotted the distribution of *GATA2* RNA punctae per cell, we found no difference between the GATA2^low^ and GATA2^high^ groups (Fig 1J-K), implying similar mRNA levels in both groups despite highly divergent levels of GATA2 protein.

### GATA2 protein is rapidly SUMOylated, poly-ubiquitinated, and degraded by the proteasome

As transcriptional regulation failed to account for differences in GATA2 protein levels, we next examined whether post-translational mechanisms may impact USC GATA2 protein stability. We utilized USC cell lines for these studies, despite our findings suggesting that transcriptional mechanisms are likely a primary determinant of cell line GATA2 levels (Fig 1A-B). We reasoned that GATA2 post-translational mechanisms would likely also be active in USC cell lines, and mechanistic studies in primary USC tumors from patients are not feasible. To identify pathways that may regulate USC GATA2 stability, we performed a cycloheximide chase experiment in the Ark1 USC cell line and co-treated cells with either vehicle or small molecules that inhibit common post-translational pathways: C646 (protein acetyltransferase inhibitor), RO-3306 (CDK1 inhibitor), staurosporine (STS, pan-kinase inhibitor), ML792 (SUMOylation inhibitor), and MG132 (ubiquitin-dependent proteasome inhibitor). We found that GATA2 loss follows a one-phase decay model with a calculated half-life of only 128 minutes in vehicle-treated cells (Fig 2A-B). Compared to vehicle control, treatment with ML792 or MG132 significantly prolonged GATA2 half-life, while treatment with C646, RO-3306, and STS had no impact on half-life. The results suggested that SUMOylation and ubiquitin-dependent proteasome activity normally drive GATA2 degradation in USC cell lines.

**Figure 2:**
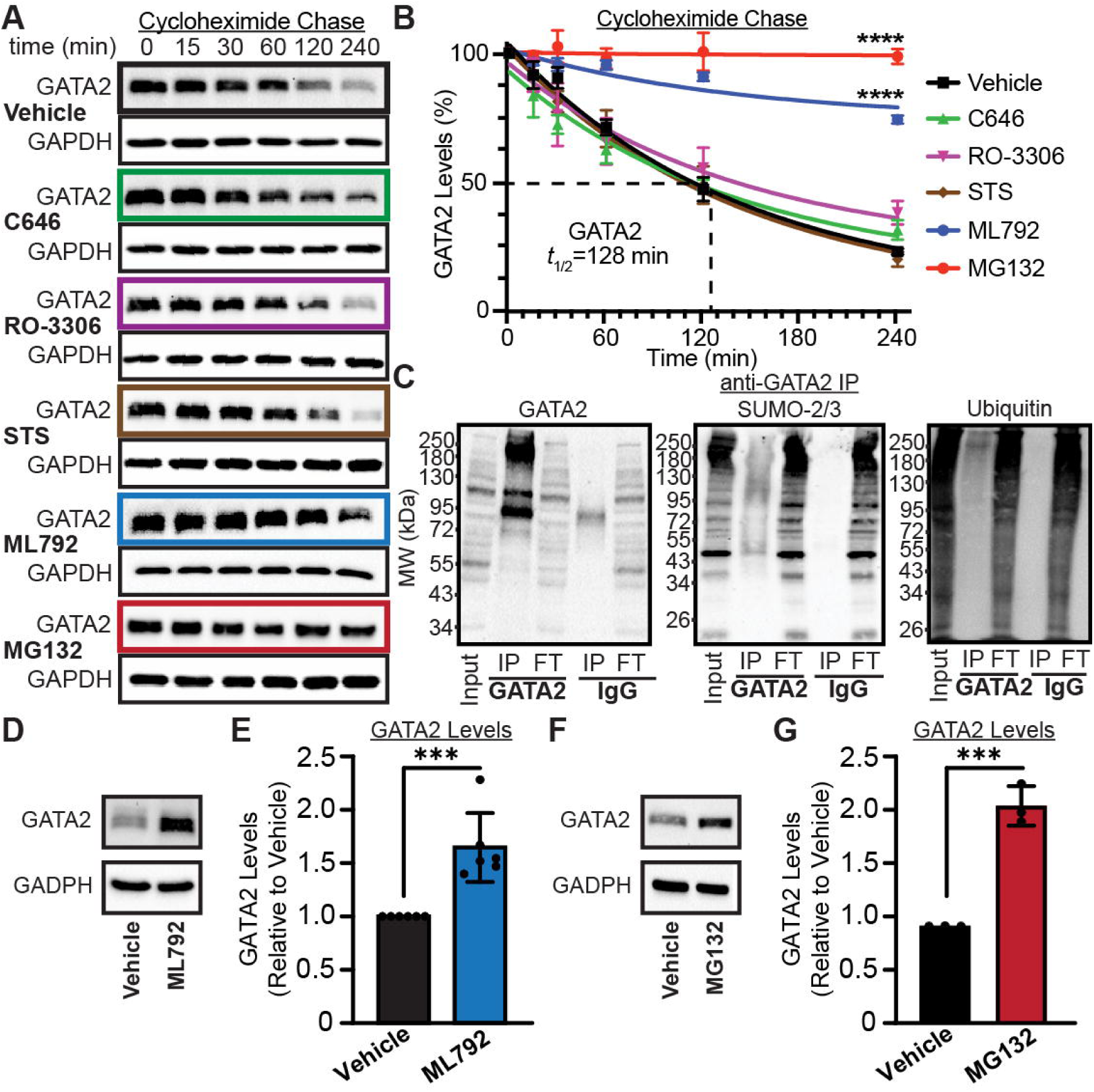
GATA2 stability is reduced by SUMOylation and ubiquitin-dependent proteome activity. **(A-B)** Cycloheximide chase quantifying GATA2 stability in Ark1 USC cells treated with vehicle, protein acetyltransferase inhibitor C646, CDK1 inhibitor RO-3306, pan-kinase inhibitor Staurosporine (STS), SUMOylation inhibitor ML792, and proteasome inhibitor MG132. **(C)** Anti-GATA2 immunoprecipitation from Ark1 USC cells and western blotting for GATA2, SUMO-2/3, and ubiquitin. IP=immunoprecipitate, FT=flow through. **(D-E)** GATA2 levels in Ark1 USC cells treated with ML792 or vehicle for 24 hours. **(F-G)** GATA2 levels in Ark1 USC cells treated with MG132 or vehicle for 8 hours. ***=p<0.0005, ****=p<0.00005.

The small Ubiquitin-like Modifier (SUMO) protein family consists of four isoforms, SUMO-1 through SUMO-4, which are attached to and detached from proteins to impact their function and/or stability.^28,29^ The SUMO-2 and SUMO-3 isoforms are nearly identical in both sequence and structure, and have largely overlapping functions and targets, and are therefore commonly denoted as SUMO-2/3.^30,31^ SUMO-1, SUMO-2, and SUMO-3 can modify overlapping substrates and acceptor lysines, but their conjugation patterns and biological effects are not identical. We chose to focus on isoforms SUMO-2/3 rather than SUMO-1 and SUMO-4 for several reasons. SUMO-1 is nearly quantitatively conjugated at steady state to constitutive substrates, leaving little free mobilizable reserve.^31^ SUMO-1 also does not form polymeric chains that can target SUMOylated proteins for degradation.^32^ SUMO-4 peptides are absent in CPTAC USC proteomes and SUMO-4 protein is not detected in benign or malignant uterine tissue or cell lines in the Human Protein Atlas.^33^ Evidence also suggests that SUMO-4 is not efficiently matured to a conjugatable form.^34^

To verify that the GATA2 protein is SUMOylated and ubiquitinated, we immunoprecipitated GATA2 from Ark1 USC cells and showed by western blot that SUMOylated and ubiquitinated species were detected in the GATA2 IP but not in the IgG control (Fig 2C). At least three SUMOylated GATA2 species were identified, ranging in molecular weight from approximately 95 kDA to 120 kDa, and 250 kDa, while the ubiquitin western blot showed only a single species at approximately 250 kDa. This is consistent with stepwise poly-SUMOylation, followed by ubiquitination. We confirmed direct GATA2 SUMOylation by performing a reciprocal anti-SUMO-2/3 IP and blotting with anti-GATA2 antibody (Suppl Fig 2).

To test whether prolonging GATA2 protein half-life by inhibiting SUMOylation and proteasome activity increases total USC GATA2 levels, we treated Ark1 cells with ML792 or MG132 and confirmed elevated total GATA2 protein levels by western blot (Fig 2D-G). ML792 or MG132 treatment also increased GATA2 levels in the unique Ark2 USC cell line, which is derived from a different patient tumor than Ark1 (Suppl Fig 3A-C). GATA2 levels also increased following siRNA-mediated SUMO-2/3 depletion (Suppl Fig 3D-F).

### A GATA2 K222R/K389R SUMOylation mutant exhibits a prolonged half-life and exhibits elevated steady-state expression levels relative to wildtype GATA2

As our data indicated that GATA2 is directly SUMOylated by SUMO-2/3 and SUMOylation activity promotes GATA2 degradation, we next sought to identify the target amino acids within GATA2 that bind SUMO-2/3. The SUMO-2/3 isoforms are considered “stress-responsive” SUMO proteins that exist mostly as a free pool. In this form, they are readily mobilized in response to cellular stimuli,^31,35^ although SUMO-2 can also modify a small number of substrates in the steady state. The latter basal activity may help to explain why *Sumo2*-null mice die in mid-gestation, whereas *Sumo3*-null mice are viable.^36^ In the laboratory, induction of the stress response is classically accomplished by the introduction of heat shock, which is known to redistribute SUMO-2/3 to hundreds or even thousands of substrates.^35,37,38^ Under stress response conditions, SUMO-2/3 is responsible for the majority of SUMOylation sites in human cells.

We first mined a large database of stress-responsive SUMOylated residues, which identified GATA2 K389 as being uniquely SUMOylated after heat shock.^37^ In addition, we noted a report of GATA2 SUMOylation in the mouse vascular endothelium, which implicated both GATA2 K222 and K389 as being SUMOylated in graft arteriosclerosis.^39^ We generated over-expression constructs for wildtype GATA2, GATA2^K222R^, GATA2^K389R^, and GATA2^K222R+K389R^ (Fig 3A). We then generated GATA2 knock-out (GATA2^KO^) Ark1 cells using CRISPR/Cas9 (Suppl Fig 4), which allowed us to overexpress wildtype GATA2 and GATA2 SUMOylation mutants in the absence of endogenous protein.

**Figure 3:**
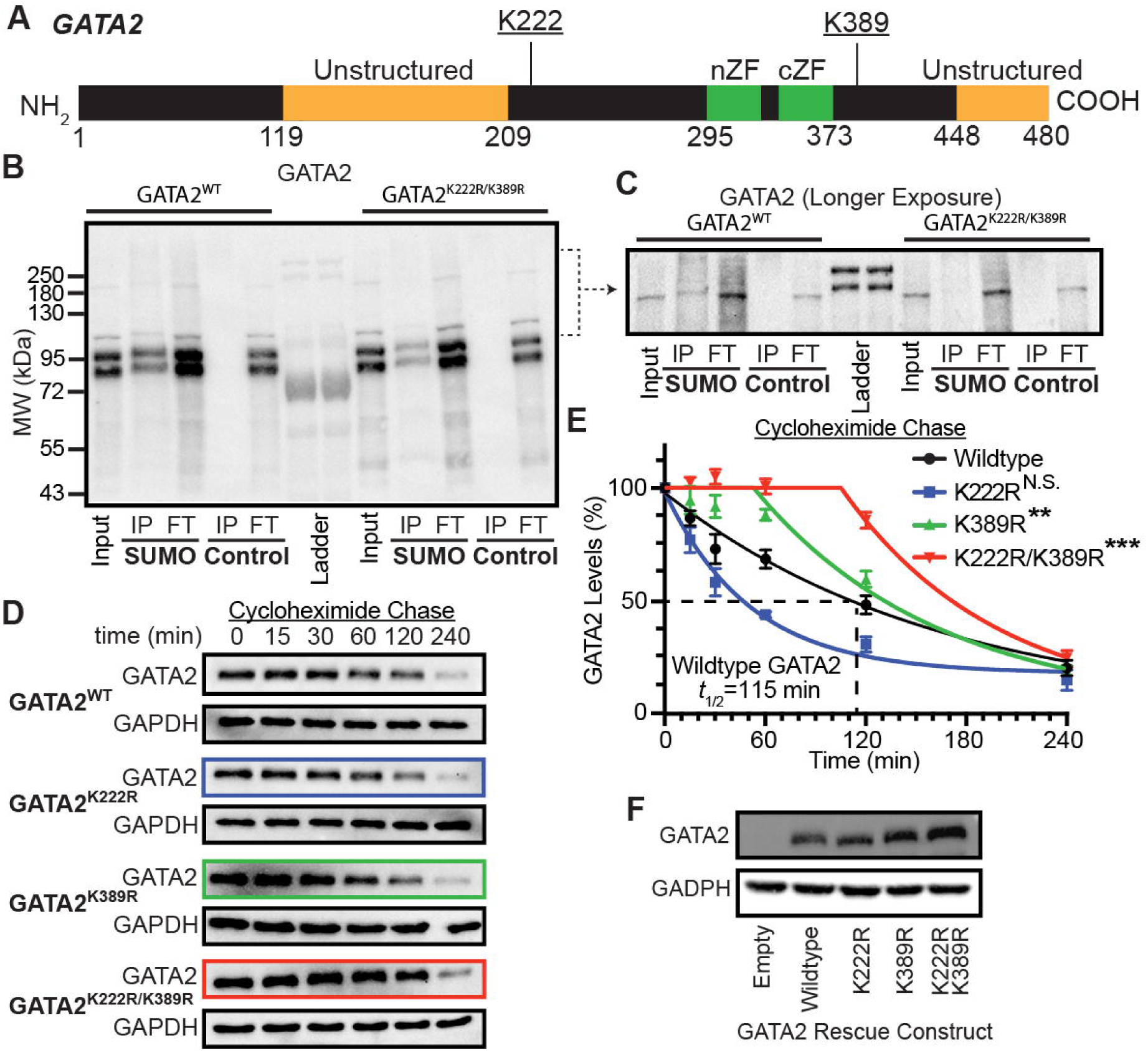
GATA2 is destabilized by K222 and K389 SUMOylation. **(A)** Domain structure of GATA2 as annotated by UniProt. Two zinc finger binding domains termed the N-terminal zinc finger (nZF) and C-terminal zinc finger (cZF) are highly conserved within the GATA factor family. These are bookended by two unstructured domains. Location of proposed K222 and K389 SUMOylation sites are also indicated. **(B)** Anti-GATA2 western blot of SUMO-2/3 immunoprecipitation (IP) from GATA2 knock-out (GATA2^KO^) Ark1 USC cells over-expressing wildtype GATA2 (GATA2^WT^) or GATA2^K222R+K389R^ mutant. **(C)** Prolonged exposure of higher molecular weight regions from panel B. **(D-E)** Anti-GATA2 western blot of cycloheximide chase experiment comparing protein stability of wildtype GATA2 and GATA2 SUMOylation site mutants overexpressed in GATA2^KO^ Ark1 USC cells. **(F)** Anti-GATA2 western blot showing total GATA2 levels of wildtype GATA2 and GATA2 SUMOylation point mutants in GATA2^KO^ Ark1 USC cells. ND=no statistical difference, **p<0.005, ***p<0.0005.

To test whether the K222R and/or K389R mutation impacted GATA2 SUMOylation, we performed anti-SUMO-2/3 immunoprecipitations followed by anti-GATA2 western blots using an antibody generated against the N-terminal 204 amino acids of GATA2^40^ that does not include the mutated residues. Similar levels of total mutant GATA2 and wildtype GATA2 were expressed (input lanes in Figure 3B and Suppl Fig 5), but a reduced GATA2 signal was detected in the SUMO-2/3 IP lane in GATA2 SUMO mutants, indicating less but not absent GATA2 SUMOylation. This was especially pronounced in the GATA2^K222R+K389R^ double-mutant, which also showed nearly complete loss of the higher molecular weight GATA2 species at 250 kDa corresponding to ubiquitinated GATA2 (SUMO IP lanes in Fig 3C).

We next tested whether mutations of GATA2 SUMOylation sites impacted GATA2 half-life. Overexpressed wildtype GATA2 had a half-life of 115 minutes, which was similar to the 128 minutes measured for endogenous GATA2 (Fig 3D-E). Interestingly, the GATA2^K222R^ mutant trended towards a shortened half-life versus wildtype control, although this was not statistically significant. However, the GATA2^K389R^ and GATA2^K222R+K389R^ mutants adopted decay kinetics that were best fit by a plateau followed by one-phase decay model, and had significantly longer half-lives that wildtype control. This resulted in higher steady state levels of the GATA2^K389R^ and GATA2^K222R+K389R^ mutants versus either the GATA2^K222R^ mutant or wildtype control (Fig 3F).

### GATA2 stabilization suppresses USC invasion

Reduced GATA2 protein levels in patient USC tumors correlate with invasion, and GATA2 depletion promotes USC invasion *in vitro*.^20^ If inhibition of GATA2 SUMOylation stabilizes GATA2, we hypothesized that the resulting increased GATA2 levels may suppress USC invasion. We first confirmed that GATA2^KO^ USC cells showed increased invasion versus wildtype controls (Fig 4A). We next rescued GATA2 expression in GATA2^KO^ cells by over-expressing wildtype GATA2 or GATA2^K222R+K389R^ mutant and tested their ability to suppress USC invasion versus empty vector control (Fig 4B). As expected, wildtype GATA2 significantly suppressed USC invasion versus empty vector control, and invasion was further suppressed by expression of the GATA2^K222R+K389R^ mutant.

**Figure 4:**
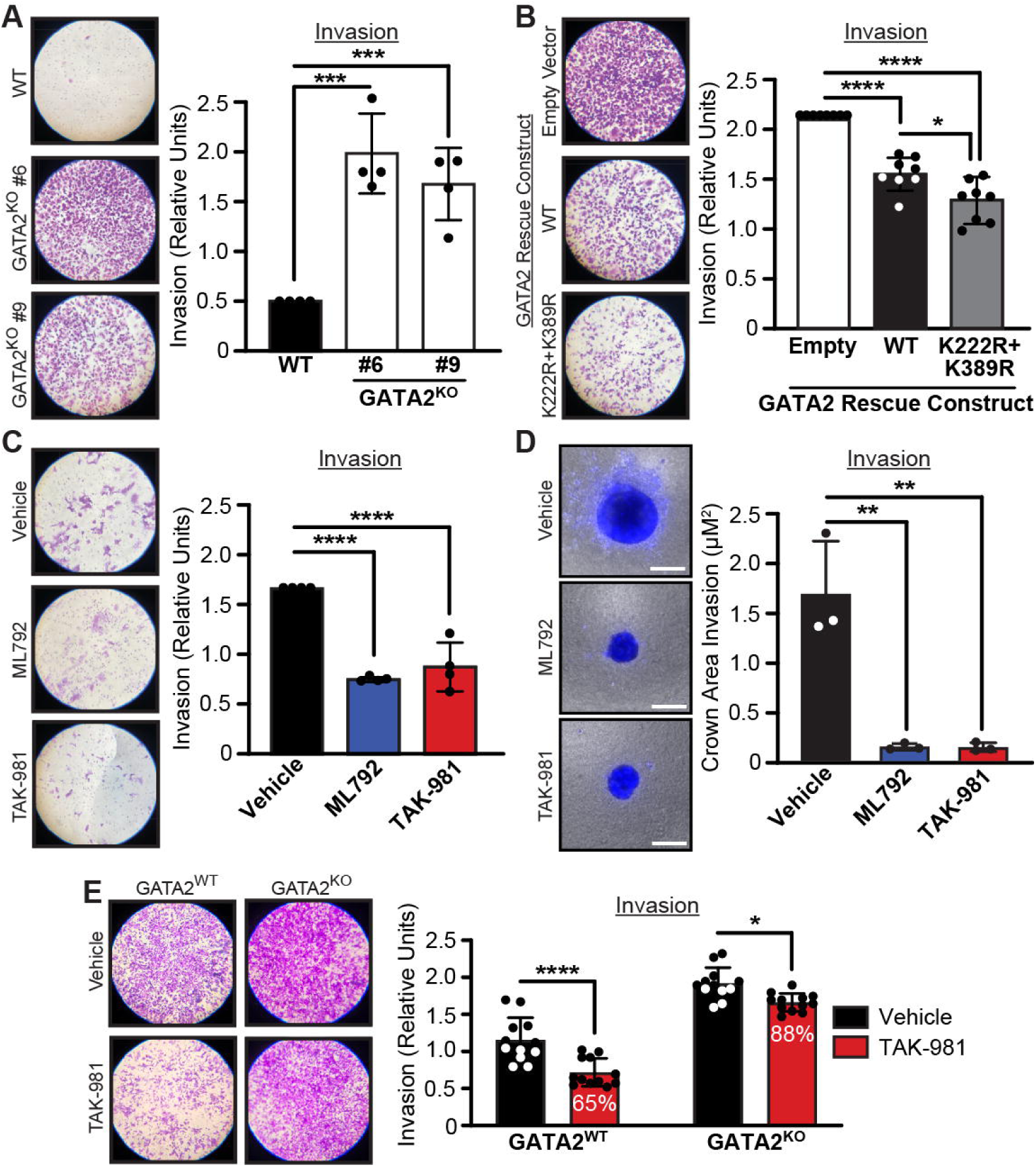
Inhibition of GATA2 SUMOylation suppresses USC invasion *in vitro*. **(A)** Invasion of wildtype (WT) Ark1 USC cells and 2 GATA2 knock-out (GATA2^KO^) clones. **(B)** Invasion of GATA2^KO^ Ark1 USC cells rescued with WT GATA2 or GATA2^K222R+K389R^ mutant. **(C)** Invasion of Ark1 USC cells treated with vehicle, ML792, or TAK-981 for 24 hours. **(D)** Invasion of Ark1 USC tumor spheroids embedded in 3D hydrogel, treated with vehicle, ML792, or TAK-981, and allowed to invade over 5 days. **(E)** Invasion of wildtype or GATA2^KO^ Ark1 USC cells treated with TAK-981 or vehicle control. Percentages in white font indicate percent reduction from vehicle. *p<0.05, **p<0.005, ***p<0.0005, ****p<0.00005.

As inhibition of SUMOylation stabilized GATA2 and increased steady-state GATA2 levels in USC cells, we tested whether SUMOylation inhibitors could suppress USC invasion. In addition to ML792, TAK-981 is a SUMOylation inhibitor that is structurally unique from ML792 and has been evaluated in Phase 1 and 2 clinical trials for solid and hematologic malignancies.^41,42^ To ensure any impact of ML792 or TAK-981 on invasion was not confounded by cell viability, we initially evaluated whether treatment with ML792 or TAK-981 impacted cell viability over 24 hours, which is the duration of our standard *in vitro* invasion assays. Neither ML792 nor TAK-981 significantly impacted cell viability over 24 hours compared to vehicle control (Suppl Fig 6A). However, both drugs significantly suppressed invasion of Ark1 USC cells *in vitro* (Fig 4C). Because USCs typically grow as non-invasive three-dimensional tumors prior to myometrial invasion, we sought out an additional and more physiologically relevant system to test the effects of SUMOylation inhibitors on USC invasion. To this end, we generated USC tumor spheroids in hanging droplets and then casted them within closed-system three-dimensional hydrogels that allowed us to flow in nutrients and small molecules.^43–45^ At day 0, we flowed in either vehicle, ML792, or TAK-981, and after 5 days we measured tumor invasion into the hydrogel beyond the starting spheroid core volume (Fig 4D). This revealed a striking reduction in invasion after treatment with either ML792 or TAK-981 relative to vehicle. This finding was not due exclusively to cell death, as Calcein AM staining was comparable across treatment conditions at the 5-day endpoint (Suppl Fig 6B).

As GATA2 half-life also depends on polyubiquitin-mediated degradation by the proteasome, we further tested whether treatment with the FDA-approved proteasome inhibitors Bortezomib or Ixazomib^46,47^ may suppress USC invasion. However, whereas 8 hours of treatment with these agents did increase GATA2 levels in USC cell lines compared to vehicle control (Suppl Fig 7A-D), treatment for 24 hours led to marked cellular toxicity which precluded treatment for the 1-day and 5-day periods needed to assess cell line and tumor spheroid invasion, respectively (Suppl Fig 7E-F).

SUMOylation regulates many cellular proteins and processes, and the anti-tumor effects of TAK-981 are hypothesized to include both tumor-intrinsic and tumor-extrinsic mechanisms.^42,48,49^ However, our *in vitro* systems contained only USC cells, and we hypothesized that SUMOylation inhibitors suppressed USC invasion in a cell-intrinsic manner predominantly by stabilizing GATA2 protein. To confirm that the ability of ML792 and TAK-981 to suppress USC invasion relied on increasing the abundance of GATA2 protein, we treated wildtype and GATA2^KO^ cells with TAK-981 or vehicle. We reasoned that if TAK-981 restrained USC invasion by stabilizing GATA2 levels, then its ability to suppress invasion would be significantly reduced or absent in GATA2^KO^ cells. As expected, TAK-981 suppressed invasion by wildtype GATA2 cells by 35% compared to vehicle control (Fig 4E). However, TAK-981 suppressed invasion by GATA2^KO^ cells by 12%, implying its mechanism of action depends on the presence of GATA2. This result is supported by our overexpression studies demonstrating that the GATA2^K222R+K389R^ mutant suppresses invasion more than wildtype GATA2 (Fig 4B), indicating that reduced SUMOylation of GATA2 alone is sufficient to suppress USC invasion.

### SUMOylation signaling pathways are broadly upregulated in USC

As our findings indicated that post-translational regulation primarily determines GATA2 levels in USC tumors, and we elucidated GATA2 SUMOylation as a key regulator of USC GATA2 stability and steady-state levels, we hypothesized that SUMOylation pathways are upregulated in USC given that 67% of stage I tumors show a reduction or complete loss of GATA2 protein.^20^ We therefore mined CPTAC USC proteomics datasets and compared SUMOylation levels and SUMOylation pathway activity between USC and the most common histologic subtype of uterine cancer called uterine endometrioid carcinoma (UEC), and benign endometrium.

We initially observed that levels of SUMO-1, SUMO-2, and SUMO-3 were all significantly elevated in USC compared to UEC and benign endometrium (Fig 5A-B). SUMO-4 was not included in the analysis, since we failed to detect it across the CPTAC uterine cancer dataset. We hypothesized that increased SUMO protein levels were a result of elevated SUMOylation pathway activity. We tested whether gene signatures of SUMOylation, deSUMOylation, ubiquitination, proteasome activity, and proteinase activity were upregulated in USC versus UEC using Gene Set Variation Analysis (GSVA). As a control, we also compared signatures of protein neddylation, a process related to SUMOylation in which a different ubiquitin like protein (NEDD8) is covalently attached to lysine residues of target proteins, which may then serve as a substrate for targeted poly-ubiquitination, and degradation.^50,51^ USC tumors in the CPTAC Endometrial Cancer Discovery dataset^52^ showed a significant positive enrichment for deSUMOyation compared to UEC, while neddylation showed a significant negative enrichment (Fig 5C). The Reactome SUMOylation signature showed a modest but significant positive enrichment in USC versus UEC. However, the GOBP Protein SUMOylation signature showed no significant enrichment and neither did the BioCarta SUMO Pathway signature. In contrast, we found no enrichment for signatures of poly-ubiquitination, proteasome activity, or proteinase activity. We performed the same analysis on the CPTAC Endometrial Cancer Confirmatory dataset,^53^ which confirmed the significant positive enrichment for deSUMOylation and negative enrichment for Neddylation in USC versus EUC (Supplementary Fig 8). The analysis collectively supports a differential SUMOylation activity and reduced Neddylation activity in USC relative to UEC, whereas poly-ubiquitination and proteasome activity are not significantly altered.

**Figure 5:**
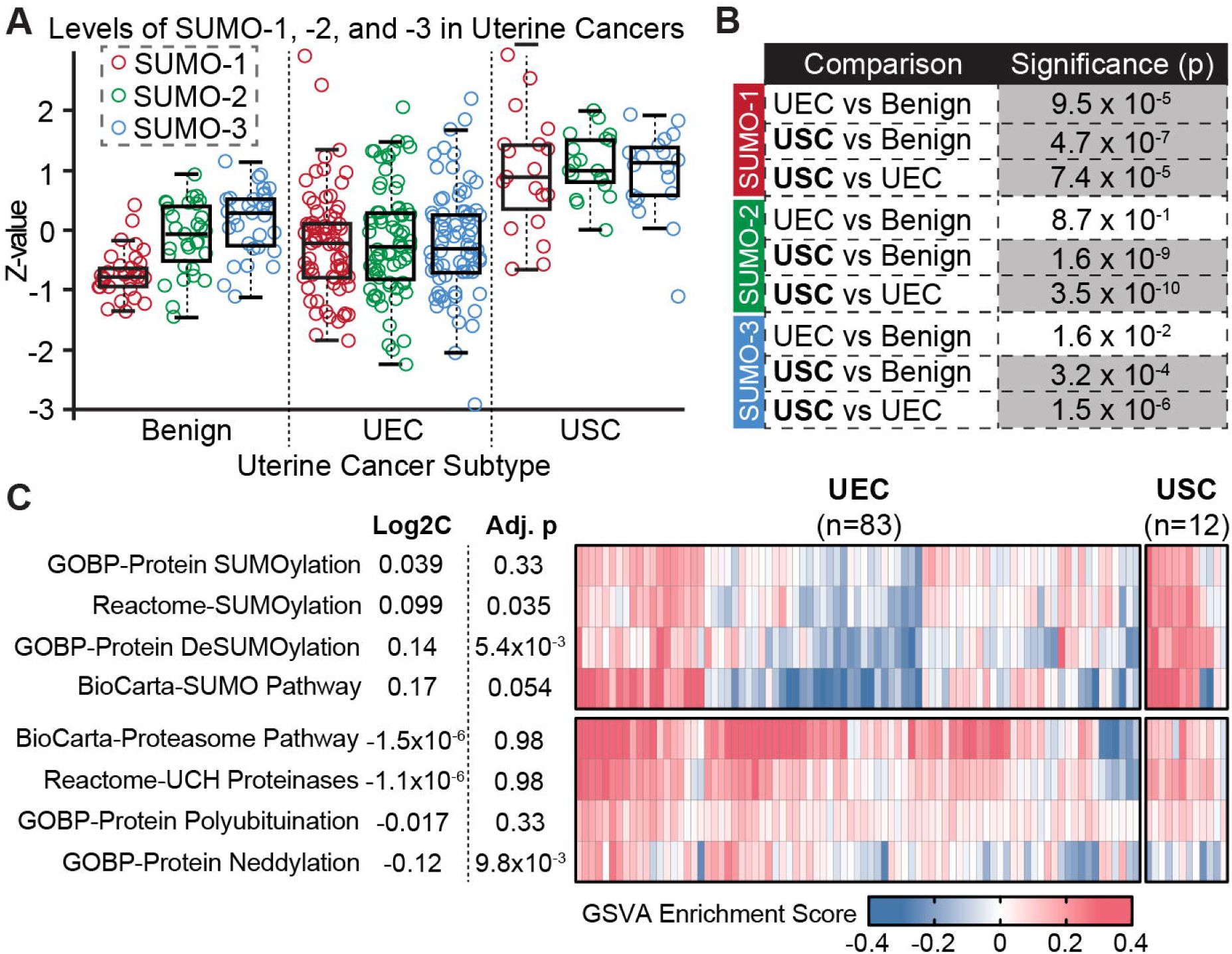
SUMOylation pathway activity is upregulated in USC versus uterine endometrioid carcinoma (UEC) and benign endometrium (benign). **(A-B)** Levels of SUMO-1, SUMO-2, and SUMO-3 protein in USC, UEC, and benign endometrium in the Clinical Proteomic Tumor Analysis Consortium (CPTAC) database. All SUMO paralogs are significantly increased in USC versus UEC and benign. Grayed out boxes indicate statistical significance. **(C)** Results of gene set variation analysis (GSVA) performed on the CPTAC Endometrial Cancer Discovery dataset^52^ using gene sets for SUMOylation, deSUMOylation, proteasome pathway, polyubiquitination, proteinases, and neddylation.

### Stress-induction of SUMOylation leads to rapid GATA2 depletion

Our data support a model whereby upregulation of SUMOylation pathway activity in USC tumors leads to GATA2 SUMOylation, which destabilizes the protein and leads to reduced steady-state GATA2 levels. To test this, we utilized heat shock. Acute heat shock is the most widely used and best-characterized stimulus for inducing SUMO-2/3 conjugation and is the reference condition for system-wide SUMO proteomic studies.^31,35^ We heat shocked Ark1 USC cells by placing them at 43 °C and measured the levels of SUMOylation and GATA2 over time (Fig 6A-B). To confirm that any differences in GATA2 levels were due to induction of SUMOylation and not an alternative heat shock-induced signaling pathway, we co-treated heat shocked cells with ML792, TAK-981, or vehicle. Heat shock triggered a rapid induction of SUMOylation in vehicle treated USC cells, which led to a significant depletion of GATA2 by 90 minutes. However, both increased SUMOylation and GATA2 depletion could be blocked by treatment with ML792 or TAK-981.

**Figure 6:**
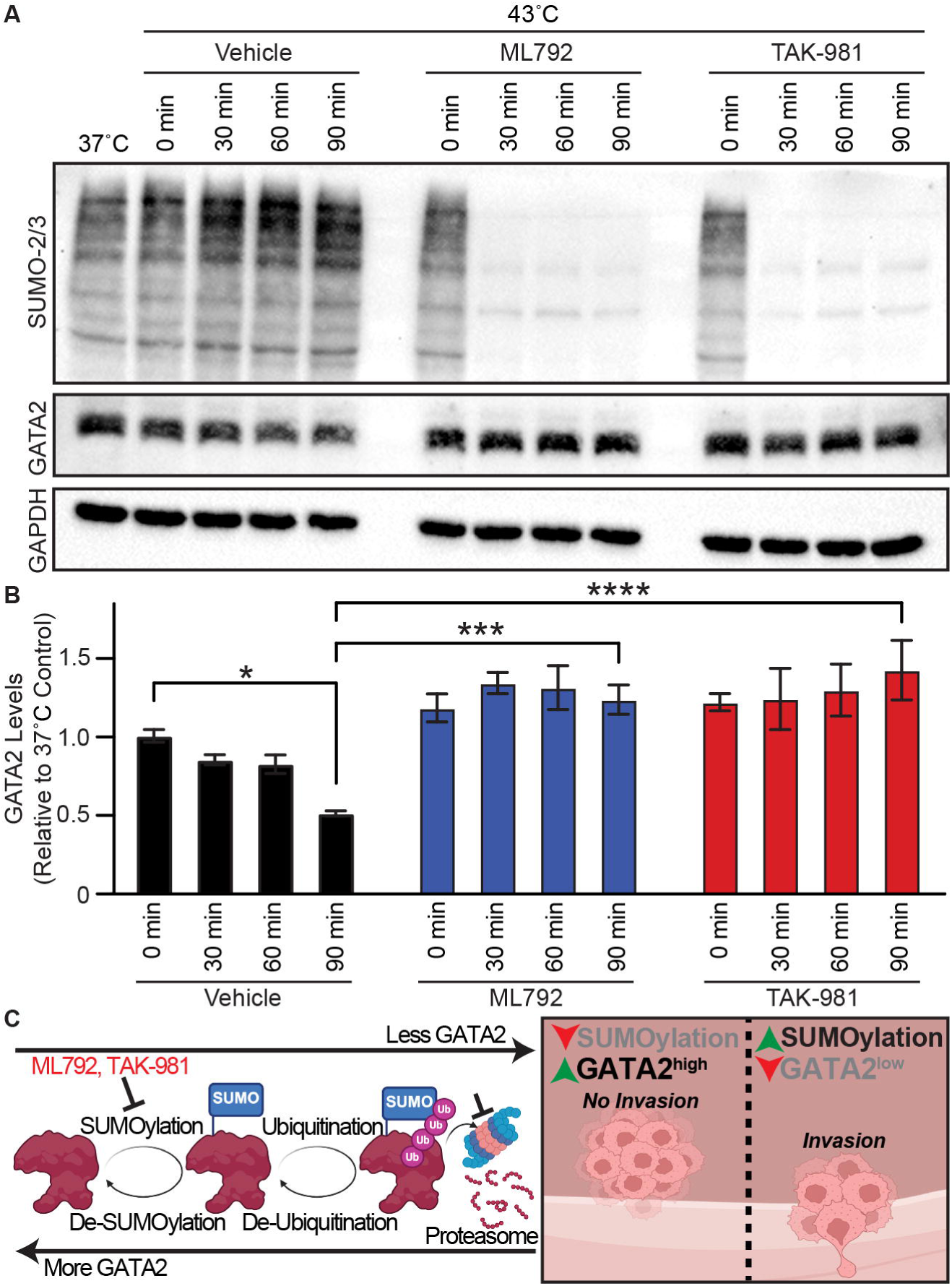
Heat shock induced SUMOylation signaling leads to SUMOylation-dependent GATA2 depletion. **(A)** Anti-SUMO-2/3 and anti-GATA2 western blot of Ark1 USC cells following induction of the heat shock response at 43°C and co-treatment with vehicle, ML792, or TAK-981. **(B)** Quantification of GATA2 levels from the heat shock experiment in panel A. **(C)** GATA2 protein stability is regulated by SUMOylation, polyubiquitination, and proteasome-mediated degradation. SUMO and ubiquitin marks can be both added and removed, while SUMOylation can be inhibited by small molecules ML792 and TAK-981. Low SUMOylation activity leads to increased GATA2 levels, which suppresses USC invasion. Induction of SUMOylation leads to reduced GATA2 levels and triggers USC invasion. *p<0.05, ***p<0.0005, ****p<0.00005.

## Discussion

We reported earlier that loss of the GATA2 protein in USC triggers myometrial invasion and predicts poor outcome for USC patients.^20^ Conversely, high GATA2 expression in USC tumors is the strongest available predictor of recurrence-free and cancer-specific survival in patients with FIGO stage I USC.^20^ These findings emphasize the importance of identifying the mechanism(s) that regulate GATA2 expression in USC tumors in order to devise targeted interventions to prevent GATA2 loss and/or restore GATA2 expression.

The present study provides initial evidence for SUMO-dependent proteolysis as a major regulatory pathway of GATA2 expression in USC cells (Fig 6C). We show that USC GATA2 is an unusually short-lived protein, with a half-life of 128 minutes, and its turnover is governed by SUMO-2/3 conjugation that licenses ubiquitin-dependent proteasomal degradation. Our study provides three independent lines of evidence to establish SUMOylation as a key determinant of GATA2 levels: 1) pharmacologic inhibition of SUMOylation and siRNA-mediated depletion of SUMO-2/3 in human USC cell lines increase stead-state GATA2 expression; 2) mutations of the SUMO acceptor lysines K222 and K389 of GATA2 prolongs GATA2 half-life in USC and elevate steady-state levels; and 3) acute induction of SUMO signaling depletes GATA2 in USC within 90 minutes, an event that is prevented by two structurally distinct SUMOylation inhibitors.

Our study recognized that the key regulators of GATA2 levels in patient USC tumors differ from those in human USC cell lines, with the latter being the primary model for studying protein turnover. GATA2 RNA and protein expression correlate tightly in USC cell lines, implying transcriptional regulation of GATA2 protein levels. A tight coupling of GATA2 RNA and protein levels also is observed in endometriosis, gastric cancer, and acute myeloid leukemia in which *GATA2* RNA levels negatively correlate with *GATA2* gene methylation, a finding consistent with transcriptional regulation via hypermethylation-mediated gene silencing.^24–26^ However, in direct contrast, our study reveals that *GATA2* RNA and methylation levels in USC tumors fail to correlate with GATA2 protein expression, which alone predicts cancer invasion and recurrence in USC patients. The most parsimonious interpretation of these data is that transcriptional control of GATA2 remains operative in USC, but post-transcriptional/translational processes predominantly control GATA2 protein levels, thereby representing therapeutic targets to restore GATA2-mediated protection to USC patients.

We also considered that progesterone via PGR-mediated signaling may regulate GATA2 protein levels in USC. GATA2 and PGR protein levels correlate closely in USC^20^ and GATA2 and PGR closely co-regulate each other’s expression in the benign endometrium.^18,19^ Yet our findings indicate that PGR protein does not significantly correlate with GATA2 RNA in USC, arguing that the transcriptional GATA2-PGR circuit that operates in normal endometrium is not what sustains their expression in USC. More generally, these data caution that RNA is a poor surrogate for GATA2 protein in USC, with the practical consequence that transcriptome-based classifiers, including those derived from TCGA, will systematically fail to capture the GATA2 axis.

Two lysines account for much of GATA2 instability, and they are not equivalent. K389R alone, and K222R/K389R in combination, converted GATA2 decay from a simple one-phase process to a plateau followed by decay, suggesting that SUMOylation acts early to commit newly synthesized GATA2 to degradation rather than acting uniformly across the existing pool. Our anti-GATA2 and anti-SUMO-2/3 studies support this conclusion and show that virtually all GATA2 protein is SUMOylated. In contrast, K222R alone did not stabilize GATA2 and trended toward a shortened half-life, a result we cannot explain. One possibility is that the K222 mutation eliminated GATA2 interaction with a stabilizing molecule, redirecting GATA2 toward K389-dependent turnover. Both mutants retained residual SUMO-2/3 conjugation, indicating that additional acceptor sites remain to be mapped. Nonetheless, the double mutant showed near-complete loss of the 250-kDa ubiquitinated GATA2 species, placing poly-SUMOylation upstream of ubiquitin transfer and implicating a SUMO-targeted ubiquitin ligase as the intervening step. RNF4 and RNF111 are the leading ligase candidates, and they would offer a substantially more selective target than the SUMO E1. Finally, our findings are consistent with a localized abnormality in the SUMO pathway contributing to the loss of GATA2 protein in USC tumors, rather than alterations in more global protein turnover, because USC tumors were differentially enriched for SUMOylation activity signatures versus endometrioid tumors but showed no enrichment for ubiquitination, proteasome, or proteinase signatures.

We hypothesize that stress-induced SUMO pathway activity is the physiologic trigger for GATA2 depletion. We employed heat shock to provide a tractable, well-characterized means of stress induction,^35,37^ while acknowledging that heat shock is not a stress that USC encounters and the identity of the endogenous stimulus is unresolved. Hypoxia, therapy-induced stress, and the proteostatic demands of rapid proliferation are candidates for stress induction that are not mutually exclusive. Distinguishing among them, ideally *in vivo* and with SUMO pathway activity measured directly in tumor tissue, is a priority for future work.

Our findings have potential therapeutic implications. TAK-981 is a SUMOylation inhibitor evaluated in clinical trials for both solid and hematologic malignancies.^41,42^ In our studies, both TAK-981 and ML792 suppressed USC invasion at concentrations that did not affect the viability of USC cell lines in either conventional invasion assays or in a closed-system three-dimensional hydrogel model. Notably, we measured the effect of these small-molecular inhibitors to be largely GATA2-dependent; TAK-981 reduced invasive capacity be 35% in GATA2-intact cells but only by 12% in GATA2^KO^ cells, implying that approximately two-thirds of its anti-invasive activity may be attributed to GATA2. Our findings also support the utility of using USC GATA2 levels as a clinical biomarker to predict drug response to SUMOylation inhibitors. Logically, SUMOylation inhibitors would be most beneficial for patients with stage I USC tumors expressing low GATA2 (GATA2^low^ USC^20^), who comprise two-thirds of stage I cases and carry essentially all of the recurrence risk. Therapeutic interventions that restore GATA2 to these USC tumors may mitigate or prevent tumor invasion or metastasis.

We have reported that the GATA2^low^ and GATA2^high^ status of USC tumors can be accurately determined across different hospital pathology laboratories using IHC and a customized GATA2-specific monoclonal antibody.^20^ This operational simplicity further enhances the value of GATA2 as a predictive biomarker to assign SUMOylation inhibitor treatment to stage 1 USC patients. However, two caveats temper this optimism and require further consideration. First, a component of the antitumor activity of TAK-981 *in vivo* is thought to be immune-mediated and our USC tumor cell restricted systems do not include immune system components.^48^ Second, we observed that TAK-981 still significantly suppresses USC invasion in GATA2KO USC cells, implying that SUMOylation inhibitors partially target GATA2-independent pathways to exert this therapeutic effect. The apparent targeting by SUMOylation inhibitors of multiple SUMOylation events may broaden inhibitor efficacy but confound the use of GATA2 as a biomarker to predict drug response.

Several limitations of our study should be acknowledged. First, the proteogenomic analyses rest on small cohorts, comprising 14 USC tumors with matched quantitative proteomics in CPTAC and 24 USC Stage I tumors in TCGA. These analyses will require validation using larger tumor proteomic datasets as they become available. Second, mechanistic studies were performed in two patient-derived USC lines, and SUMOylation site mutants were evaluated by overexpression in a GATA2^KO^ background rather than by knock-in of mutants at the endogenous locus. Thus, it’s possible that absolute stoichiometry of GATA2 and regulation of downstream targets may differ from the native setting. Third, we have not performed preclinical studies to evaluate whether restoring GATA2 to GATA2^low^ USC tumors suppresses invasion or metastasis *in vivo*, which is a necessary step before clinical translation. This is complicated by the lack of an animal model that spontaneously develops USC and the absence of a tractable animal explant model of stage I USC.

In summary, our study provides initial evidence that the SUMOylation-ubiquitination pathway rapidly degrades the GATA2 protein in USC tumor cells in which the loss of GATA2 highly correlates with clinical disease recurrence and poor survival.^20^ Restoration of GATA2 in USC GATA2^KO^ cells by a SUMOylation inhibitor in clinical development restores GATA2 within hours and suppresses the invasive phenotype that is responsible for most mortality in USC patients. These findings raise the possibility that USC patients with GATA2^low^ stage I tumors identified by routine IHC may benefit from SUMOylation inhibitors or other treatments that increase GATA2, thereby offering a promising intervention for patients with few therapeutic options.

## Materials and Methods

### IRB Approval and Patient Consent

This study was approved under University of Wisconsin - Madison IRBs #2017-0765 (PI McGregor) and #2018-1510 (PI McGregor). All tumor tissues represented residual material collected during routine clinical care and patient consent was waived.

### Cell culture and Drug Treatments

Ark1 and Ark2 patient-derived USC cells were purchased from Alessandro Santin (Yale University, New Haven, Connecticut, USA) and maintained at 37°C in a humidified incubator at 5% CO_2_ using RPMI-1640 + 10% FBS + 1% penicillin/streptomycin. For cycloheximide chase experiments, Ark1 cells were seeded at 3×10^5^ cells per well in 6-well plates one day before treatment. For steady-state protein analysis, Ark1 and Ark2 cells were treated with 500 nM ML792 or 500 nM TAK-981 for 24 hrs, or 5 μM MG132 for 8 hours.

### Cycloheximide Chase

Cells were treated with cycloheximide (CHX; Sigma-Aldrich, C4859) at 50 μg/mL to inhibit new protein synthesis. Where indicated, cells were treated with 500 nM ML792, 20 μM RO-3306, 1 μM staurosporine, 25 μM C646, 5 μM MG132, or 500 nM TAK-981. Normalized GATA2 signal was fit by nonlinear regression to one-phase exponential decay or, where indicated, to a plateau followed by one-phase decay; the preferred model was identified by the extra sum-of-squares F test, and decay curves and derived half-lives were compared against vehicle or wildtype GATA2 control by with Dunnett’s multiple comparisons test.

### Methylation Analysis

DNA methylation reads were quality trimmed using Trim Galore (v0.6.10) with cutadapt (v4.5). Trimmed reads were aligned to the human reference genome (hg19) using Bismark (v0.24.2) with Bowtie2 (v2.5.1). M-bias profiles were assessed with Bismark, and bases within bias-affected regions at read ends were excluded from downstream methylation analysis (3 bp from the R1 5′ end, 6 bp from the R2 5′ end, and 2 bp from the R2 3′ end). Cytosine methylation calls were subsequently extracted using Bismark.

### Western Blotting

For whole-cell lysates, cells were lysed in sample buffer (62.5 mM Tris-HCl, pH 6.8, 2% SDS, 10% glycerol, 100 mM DTT, and 0.008% bromophenol blue). Lysates were sonicated and boiled for 5 min. When necessary, protein concentration was determined by BCA assay using lysates prepared in NP-40-based lysis buffer (25 mM Tris-HCl, pH 7.5, 150 mM NaCl, 1% NP-40, and 1 mM EDTA supplemented with protease inhibitors). Protein concentration was determined by BCA assay. The following antibodies were utilized: anti-GATA2 (15D2 mouse monoclonal), anti-GATA2 (rabbit polyclonal),^40^ anti-SUMO-2/3 (Cell Signaling Technology, #14066), anti-ubiquitin (Cell Signaling Technology, #3936S), anti-GAPDH (Cell Signaling Technology, #3683), and anti-mCherry (Cell Signaling Technology, #43590). Detection was accomplished with HRP-conjugated secondary antibodies (Jackson ImmunoResearch) and either standard ECL (ThermoFisher, #32109) or ECL pico (ThermoFisher, #PI34577). Immunoblot signals were quantified by densitometry in ImageJ, normalized to load, and expressed relative to the vehicle or empty-vector control of the same replicate.

### Immunoprecipitations

For anti-GATA2 IPs, lysates were prepared in modified 2×Laemmli buffer (150 mM Tris-HCl, pH 7.2, 4% SDS, 20% glycerol, and 20 mM freshly prepared N-ethylmaleimide (NEM)). Lysates were sonicated, boiled at 100°C for 10 min, and clarified by high-speed centrifugation for 15 min at room temperature. SDS was removed using detergent-removal spin columns (ThermoScientific, 87778/87779) according to the manufacturer’s instructions. SDS free lysates were incubated with 2 µg anti-GATA2 antibody or species-matched control IgG for 24 hrs at 4°C. Immune complexes were captured with Protein A/G magnetic beads for 24 hrs, washed four times wash buffer (50mM Tris-HCl pH 7.5, 150 mM NaCl, 1% NP-40) and eluted in SDS sample buffer.

For anti-SUMO-2/3 IPs, cells were lysed in modified 2×Laemmli buffer as previously described. Clarified lysates containing 0.5 mg total protein were incubated with SUMO-2/3 affinity beads (Cytoskeleton, ASM24-Beads) or control agarose beads for 24 hrs at 4°C with rotation. Beads were washed 4 times with SUMO IP wash buffer (50mM Tris-HCl pH 7.5, 150 mM NaCl, 1% NP-40) and bound proteins were eluted in SDS sample buffer.

### Invasion Assays

For Matrigel-based invasion assays, we utilized Matrigel-coated membrane inserts with 8.0 μm pores (Neta Scientific, #354480). Inserts were equilibrated to room temperature and rehydrated in serum-free medium before use. Ark1 cells were suspended in low-serum medium and seeded into the upper chamber, with complete medium containing serum added to the lower chamber as a chemoattractant. Cells were allowed to invade for 24 hrs at 37°C. Non-invading cells were scrubbed from the upper surface of the membrane with cotton swabs. Invaded cells on the lower surface were fixed and stained with 0.2% crystal violet prepared in buffered formalin (1 g crystal violet (Sigma-Aldrich, C0775) per 500 mL buffered formalin (Sigma-Aldrich, HT501128)). Crystal violet was destained from the stained Matrigel-coated inserts using a 1:1 mixture of water:methanol. Absorbance was measured at 570 nm and values were normalized to indicated control.

For spheroid invasion assays into 3D hydrogel, spheroids (2,500 cells/spheroid) were generated using the hanging drop method for previously-generated Ark1 shScramble cells.^20^ Briefly, cells were suspended in relevant media containing 20% methylcellulose (4,000 cP viscosity, dissolved in RMPI serum-free media) (Sigma-Aldrich, #M0512).^58,59^ 25 µL droplets of cell suspension at the relevant concentration were pipetted on the inner face of a Petri dish lid and distilled water was added to the base of the dish to reduce evaporation during spheroid formation. After 3 days, the spheroids formed in each droplet were collected using a wide orifice tip into a single tube and immediately used for subsequent hydrogel invasion assays.

A collagen hydrogel mixture was made on ice with a final concentration of 2.5 mg/mL rat-tail collagen type 1 (Corning, #354249), PBS10X (Invitrogen, #AM9624) was used to adjust osmolarity, and 0.5 M NaOH (Sigma-Aldrich, #567530, in distilled water) to adjust pH. This collagen mixture was diluted with fibronectin (Sigma-Aldrich, #F1141) and 1.28 mg/mL Matrigel (Sigma-Aldrich, #CLS356234) to achieve final concentrations of 30 µg/mL and 1.28 mg/mL, respectively.^20,60,61^ Using a wide orifice tip (Fisherbrand, #02-707-464), collected spheroids were added into the collagen mixture (30 spheroids in 50 µL) and a 7 µL droplet with a singular spheroid was pipetted into the center of a well of an optical bottom 96-well plate.^62^ Collagen hydrogels with Ark1 spheroids were polymerized upside down for 15 min at room temperature and then in a standard incubator for 30 min. After collagen polymerization, 150 µL of media containing 1 µg/mL doxycycline and 500 nM SUMOylation inhibitor (ML792 or TAK-981, resuspended in DMSO at a stock concentration of 10 mM) was added to each well. Spheroids were imaged at 0 hrs with a wide-field epifluorescence microscope (10X in phase contrast, Leica Thunder 3D imager) and then incubated for 120 hrs. Media containing drugs was replenished at 72 hrs. At 120 hrs of culture, a staining solution containing 1:2000 Hoechst 33342 in PBS (ThermoFisher, #H21486) was added to each well and incubated for 30 min. Following, Ark1 spheroids were imaged with the Leica Thunder 3D imager (10X, z stacks in phase contrast and LED excited at 361nm and emission filter of 486nm, steps of 10 µm). Stacks were z-projected using ImageJ and then the area occupied by the spheroid (core) and cells invaded (crown) was manually traced to quantify the area of invasion.

### Generation of GATA2 Knock-Out and Mutant Rescue Cell Lines

To generate GATA2 knock-out (GATA2^KO^) Ark1 cells, a synthetic crRNA targeting exon 2 of human *GATA2* (5′-ATGCCAACCCCGCTCACGCG-3′; Integrated DNA Technologies) was duplexed with ATTO647-labeled (IDT, #10007853) tracrRNA for 30 min at 37°C. The duplexed gRNA was then incubated with 1.7 µL Cas9 nuclease (IDT, #1081058) for 15 min to form an RNP complex. 3×10^5^ Ark1 cells were resuspended in 20 µL SF nucleofection solution from the Lonza SF Cell Line 4D-Nucleofector X Kit S (Lonza, #V4XC-2032), mixed with the RNP complex and 1 µL IDT electroporation enhancer, and nucleofected using program F-132 on a Lonza 4D-Nucleofector. Cells were immediately transferred to prewarmed complete medium and recovered overnight. The following day, viable ATTO 647-positive cells were isolated by fluorescence-activated cell sorting into individual wells of 96-well plates and clonally expanded. Genomic DNA was purified from expanded clones (Wizard® SV Genomic DNA Purification System, #A2361). The targeted region was amplified using forward primer 5′-GGGCTGCGTGTGCATGTTTTC-3′ and reverse primer 5′-CGCCCCTGACAGACATTGAG-3′, followed by single-end sequencing (Plasmidsaurus). FASTQ files were analyzed with CRISPResso2 v2.2.14 to quantify editing and characterize indels at individual alleles. Clones H6 and G9 were selected for further analysis. Loss of GATA2 protein was confirmed by western blotting.

The PQCXIN0-mCherry-GATA2 vector was generated using Gibson assembly and the entire sequence verified by Oxford Nanopore sequencing. K222R, K389R, and K222R/K389R substitutions were introduced by site-directed mutagenesis and verified by Sanger sequencing. The expression construct was packaged into retroviral particles produced HEK293T cells together with packaging plasmids psPAX2 and pMD2.G. Viral supernatant was added to GATA2^KO^ Ark1 cells seeded at approximately 30–50% confluence along with 10 µg/mL polybrene for 24 hrs. The media was then replaced with complete growth medium containing 500 µg/mL G418 (ThermoFisher, #AAJ63871AB) at 500 µg/mL and selection took place over 7 days. Expression of mCherry-GATA2 fusion proteins was confirmed by fluorescence microscopy and immunoblotting

### Cell Viability Assay

Cell viability was assessed in 96-well plate. Cells were treated with ML792 or TAK-981 at 100 nM, 500 nM, 1 µM, and 2 µM for the indicated period. Culture medium was replaced with 100 µL of fresh medium, and 20 µL of MTT stock solution (Sigma, #M2003) was added to each well. Background control wells contained 20 µL of MTT solution in 100 µL of cell-free culture medium. Cells were incubated at 37 °C for 4 hrs to allow formation of formazan product. Medium was replaced with 200 µL of DMSO, followed by addition of 25 µL of Sorensen’s glycine buffer (Glycine 0.1M, NaCl 0.1M, pH:10.5 by NaOH). Samples were mixed thoroughly and incubated at 37 °C for 10 min. Absorbance was measured at 540 nm.

### Heat-Shock Induction

Cells were harvested by gentle trypsinization and centrifugation. After quantifying, the cells (∼10^7^) were mixed in 700 μL of culture medium in 1 mL centrifuge tubes and subjected to heat shock at 43 °C on heat block.

### GATA2 In Situ Hybridization

*In Situ* Hybridization (ISH) was performed by the UW-Madison Translational Research in Pathology (TRIP) laboratory, using the RNAscope 2.5 HD Brown kit (BioTechne, #322300) and the RNAscope Probe Hs-GATA2 (BioTechne, #414361). Staining was performed using the FFPE standard protocol (BioTechne, Document Nos. 322452-USM and 322310-USM) with the following minor adjustments: target retrieval was performed in a steamer for 15 min, and counterstaining was performed with 100% Gill’s hematoxylin for 30 sec. Cell segmentation and ISH punctae calling were performed in QuPath.^27^

### Gene Set Variation Analyses

Global proteomics data were obtained from CPTAC UCEC discovery and confirmatory studies. Only samples with both metadata and proteomics data were retained. Cases flagged as excluded by the original publications were removed. Single-sample gene set scoring was performed with the ssGSEA method of the GSVA package (v2.4.9), consistent with the approach reported in both the Dou et al. 2020 (doi:10.1016/j.cell.2020.01.026) and Dou et al. 2023 (doi:10.1016/j.ccell.2023.07.007) papers. Four SUMOylation-related gene sets and four control gene sets related to the ubiquitin-proteasome system and post-translational modifications were evaluated. All gene sets were obtained from the Molecular Signatures Database (v2026.1). GSVA enrichment scores were compared between histologic types (endometrioid vs. serous) with linear models implemented in the limma package (v3.66.0). Surrogate variables estimated with the sva package (v3.58.0) were included as covariates in the design matrix. The robust empirical Bayes procedure was used with trend based on gene set size. Multiple-testing correction was performed by the Benjamini–Hochberg procedure.

### Statistical Analyses

Correlations were performed using Pearson (r) with a two-tailed p value. Comparisons between two groups were performed using Student’s t-test. Survival analyses were performed using Kaplan-Meier. GATA2 ISH distributions were compared using Kolmogorov-Smirnov test. Significance was assessed at p<0.05 unless otherwise stated. Invasion assays comparing three groups were analyzed by one-way ANOVA with Tukey’s multiple comparisons test, and experiments with two crossed factors, namely GATA2 genotype by TAK-981 treatment and heat-shock duration by drug treatment were analyzed by two-way ANOVA with Sidak’s correction. Analyses were performed in GraphPad PRISM.

## Data Availability Statement

The data generated in this study are available within the article and its supplementary data files.

## Acknowledgements

The authors wish to thank the University of Wisconsin-Madison Translational Research Initiatives in Pathology Laboratory (TRIPath Core Facility, RRID:SCR_028868), supported by the UW Department of Pathology and Laboratory Medicine, UWCCC (P30 CA014520), UW OVCR/WARF and the Office of The Director-NIH (S10 OD038136, S10 OD023526), for the use of its facilities, Leica Aperio AT2 slide imager, and *in situ* hybridization services. This work was supported by a grant from the V Foundation and the Wisconsin Partnership Program. AAM received support from the UW-Madison Science and Medicine Graduate Research Scholars (SciMed GRS) program and the Graduate Training in Molecular and Cellular Pharmacology Grant (T32 GM141013). MVM acknowledges University of Wisconsin Centennial Scholars Program funding, Building Interdisciplinary Research Careers in Women’s Health (BIRCWH) Training Award (NIAMS 2K12AR084227), and a kind gifts from the Diane Lindstrom Fund.

## Supplementary Figure Legends

**Supplementary Figure 1: GATA2 protein correlates with *PGR* RNA transcripts in USC. (A)** PGR and GATA2 protein show a significant positive correlation in benign endometrium from the Clinical Proteomic Tumor Analysis Consortium (CPTAC) database. **(B)** PGR and GATA2 protein also show a significant positive correlation in CPTAC USC cases. **(C)** *GATA2* RNA does not correlate with PGR protein in USC CPTAC cases, but (D) *PGR* RNA does show a close positive correlation with GATA2 protein.

**Supplementary Figure 2: GATA2 is directly SUMOylated by SUMO-2/3.** Immunoprecipitation (IP) of SUMO-2/3 from Ark1 USC cells identifies GATA2 and SUMO-2/3 by western blot relative to IgG controls. FT=flow through.

**Supplementary Figure 3: SUMOylation and proteasome activity drive GATA2 degradation. (A-B)** GATA2 levels increase in Ark2 USC cells treated with SUMOylation inhibitor ML792 for 8 hours. **(C-D)** GATA2 levels increase in Ark2 USC cells treated with MG132 for 8 hours. **(E-F)** siRNA-mediated SUMO-2/3 depletion leads to increased GATA2 levels in Ark1 USC cells. *p<0.05, **p<0.005, ****p<0.00005.

**Supplementary Figure 4: GATA2 knock-out (GATA2^KO^) Ark1 USC cell lines. (A)** CRISPR/Cas9 targeting *GATA2* exon 2 created biallelic insertion-deletions (indels) and frameshifts, resulting in **(B)** knock-out of GATA2 in 2 independent clones.

**Supplementary Figure 5: GATA2 knock-out (GATA2^KO^) Ark1 USC cell lines.** Anti-GATA2 western blots of SUMO-2/3 immunoprecipitation (IP) from GATA2 knock-out (GATA2^KO^) Ark1 USC cells over-expressing wildtype GATA2 (GATA2^WT^), GATA2^K222R^ mutant, GATA2^K389R^ mutant, or GATA2^K222R+K389R^ mutant.

**Supplementary Figure 6: Ark1 USC cell viability following treatment with SUMOylation inhibitors. (A)** Treatment of Ark1 USC cells with ML792 or TAK-981 for 24 hours has no impact on cell viability versus vehicle controls. **(B)** The viability marker Calcein AM shows similar levels of positivity in Ark1 USC tumor spheroid cells following 5-day treatment with ML792 or TAK-981 versus vehicle control. While tumor spheroids are significantly smaller and show less invasion after ML792 or TAK-981 treatment versus vehicle, cellular regions show similar levels of Calcein AM signal.

**Supplementary Figure 7: GATA2 levels in Ark1 and Ark2 cells after proteasome inhibition with Bortezomib and Ixazomib. (A-D)** Treatment of **(A-B)** Ark1 or **(C-D)** Ark2 USC cells with Bortezomib or Ixazomib for 8 hours led to a significant increase in GATA2 levels as measured by western blot. Quantification in **(B)** and **(D)** reflect n=4. **(E-F)** Viability of **(E)** Ark1 and **(F)** Ark2 USC cells after 24 hours treatment with varying concentrations of Bortezomib or Ixazomib relative to vehicle-treated controls. *p<0.05, **p<0.005, ***p<0.0005.

**Supplementary Figure 8: Gene Set Variation Analysis in uterine serous carcinoma (USC) versus uterine endometrioid carcinoma (UEC).** Data derived from the CPTAC Endometrial Cancer Confirmatory dataset.^53^

