## Supplementary figures and images for "GATA2 SUMOylation Regulates GATA2 Stability to Control Uterine Serous Carcinoma Invasion"

### Supplementary Figure 1

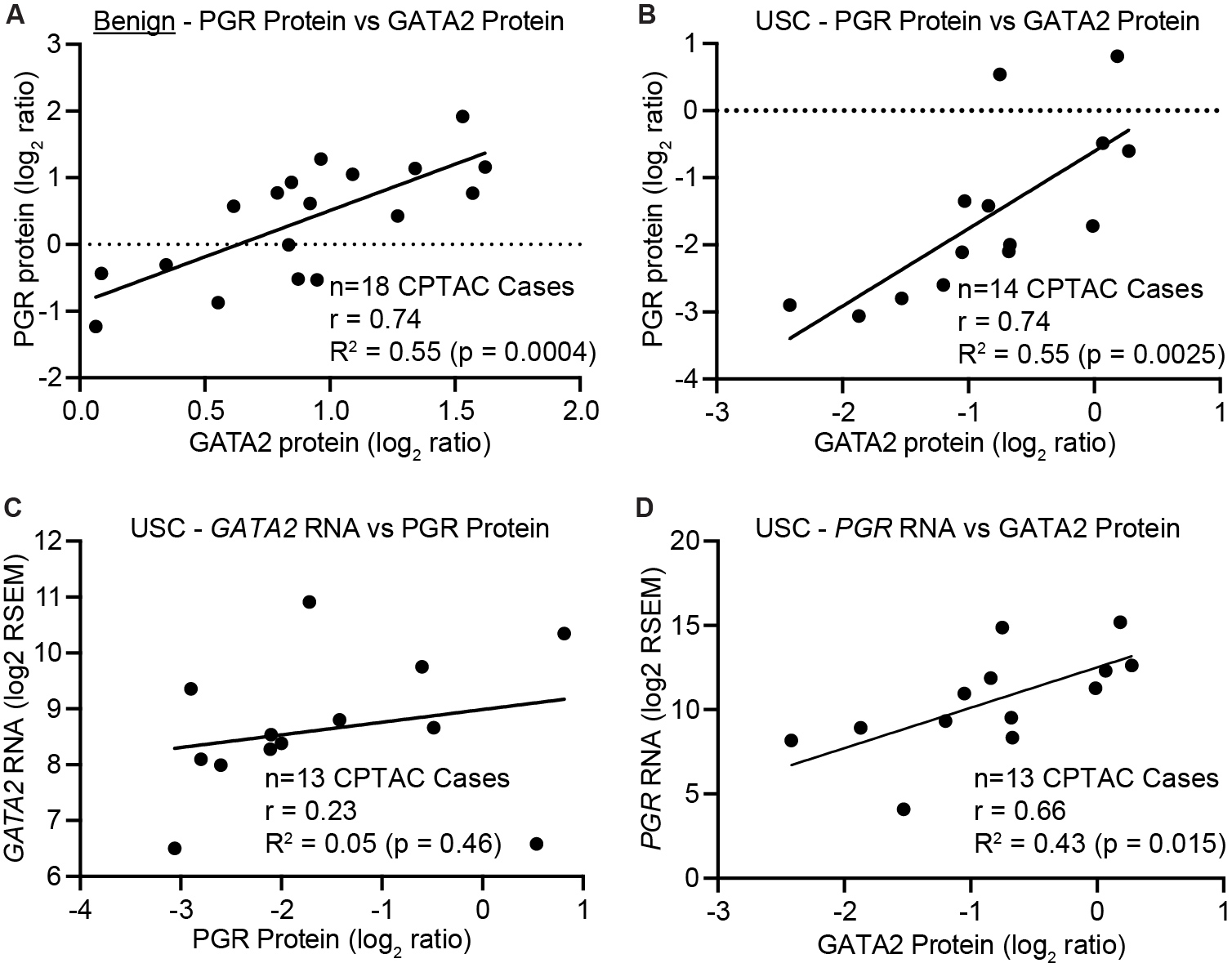

### Supplementary Figure 2

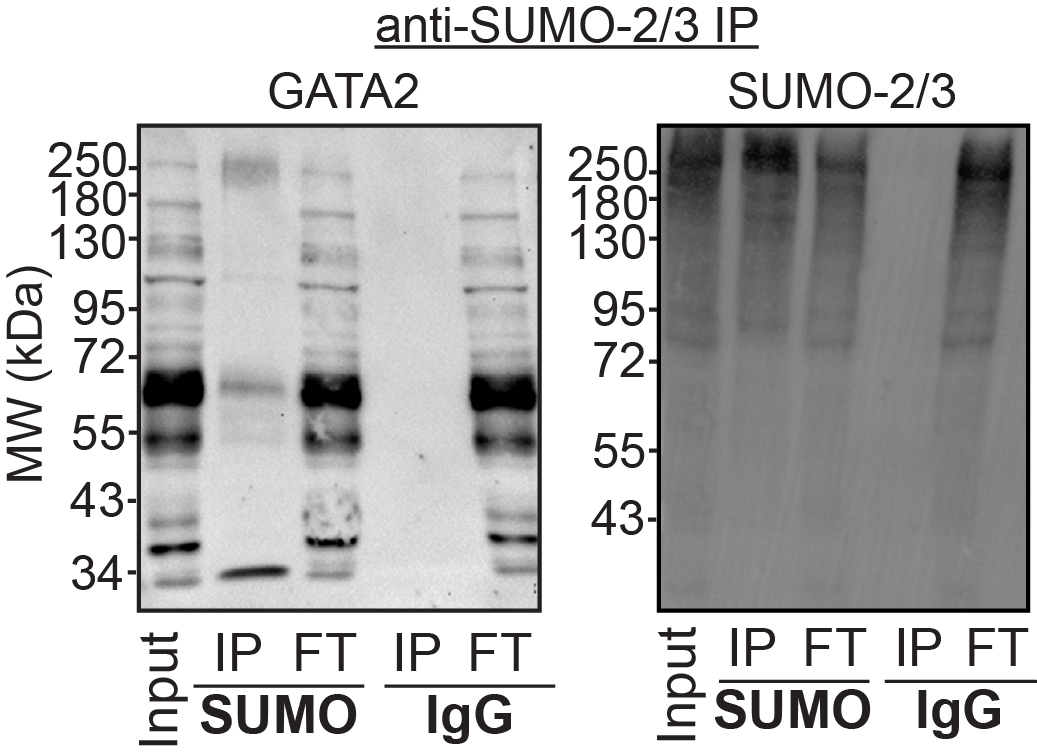

### Supplementary Figure 3

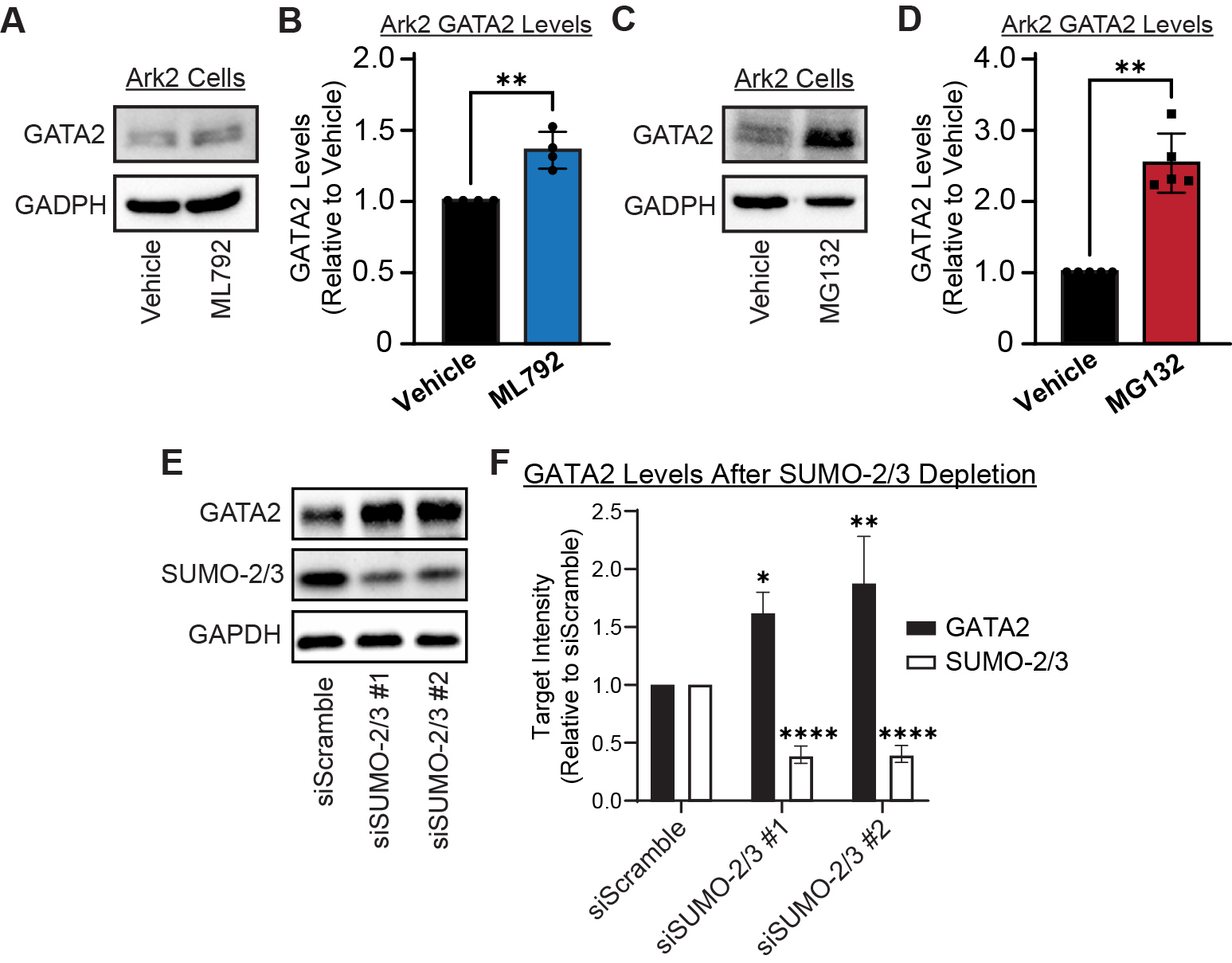

### Supplementary Figure 4

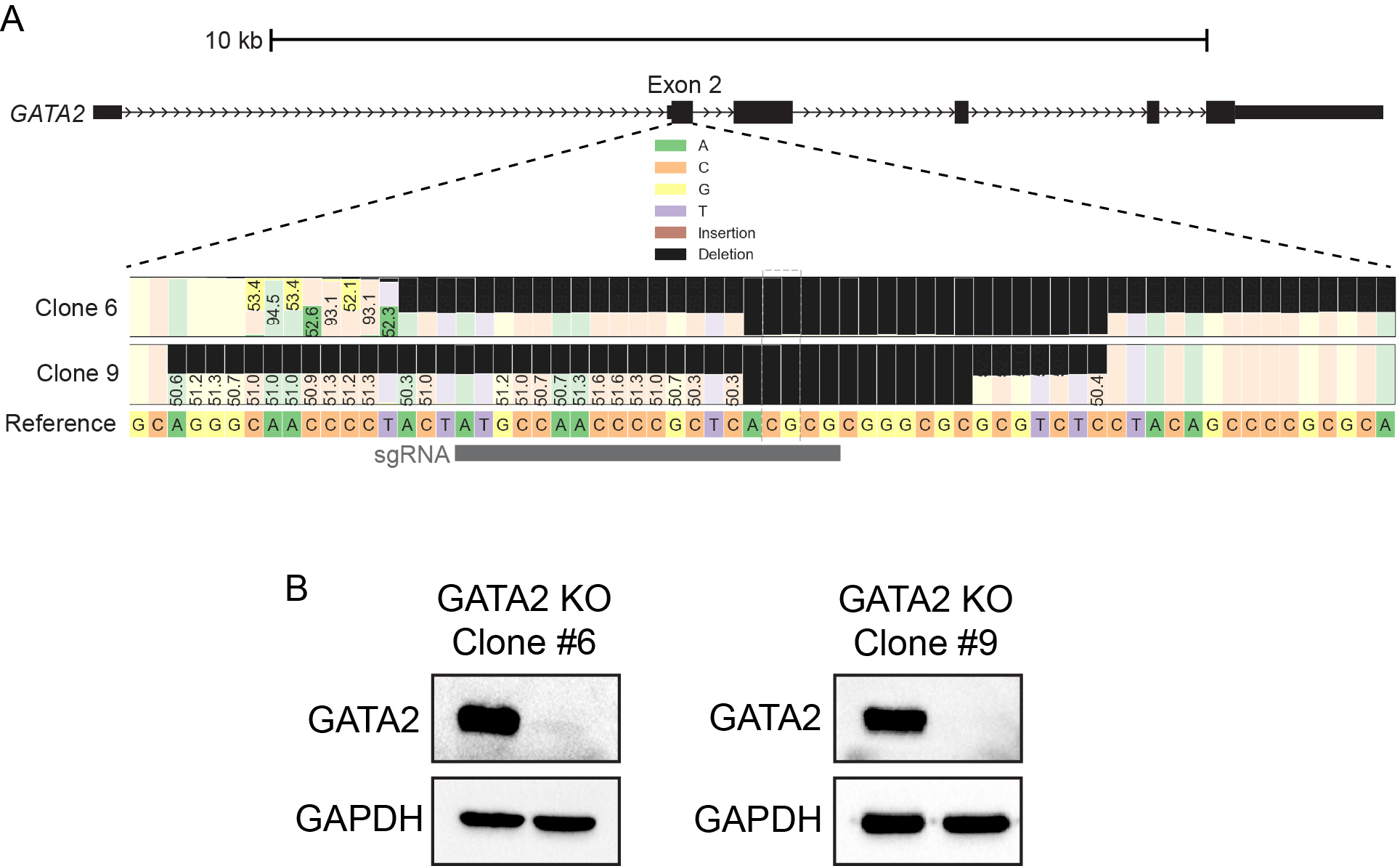

### Supplementary Figure 5

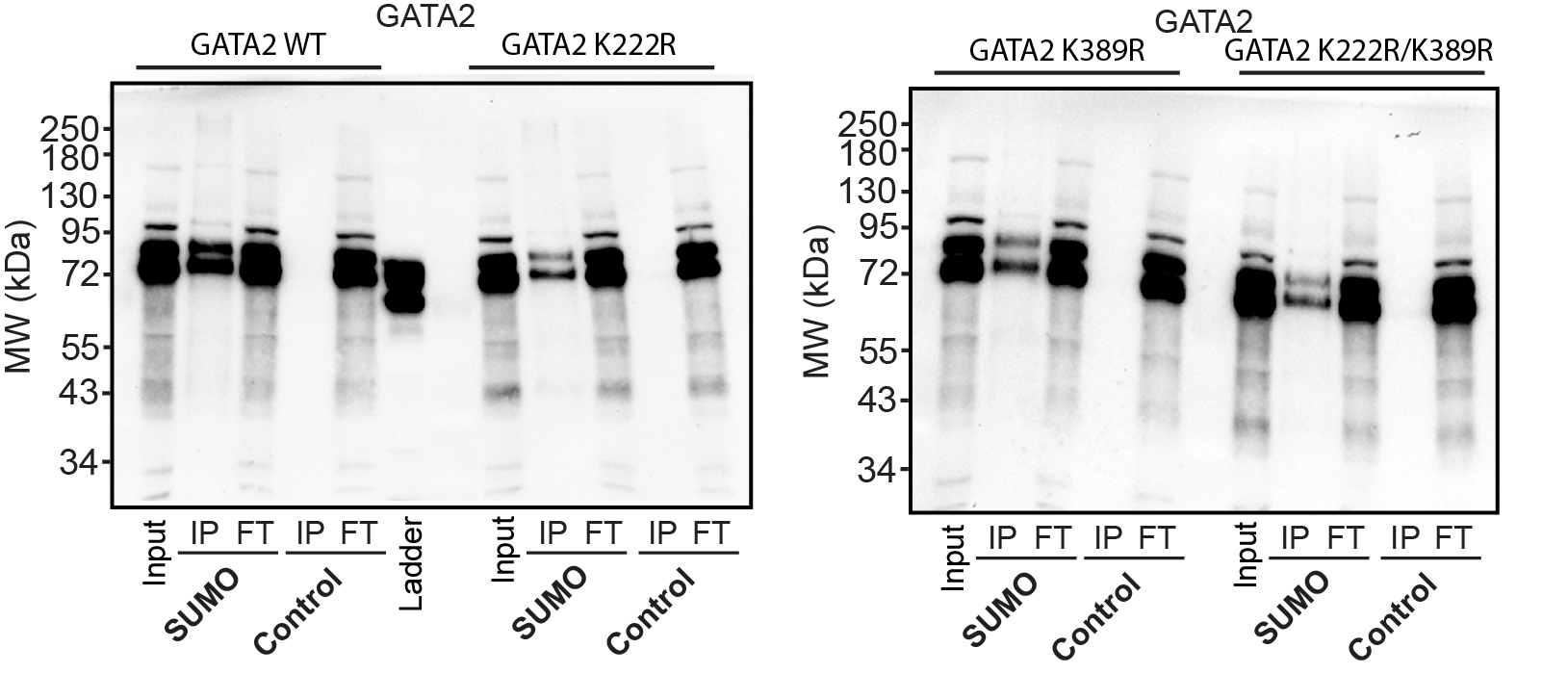

### Supplementary Figure 6

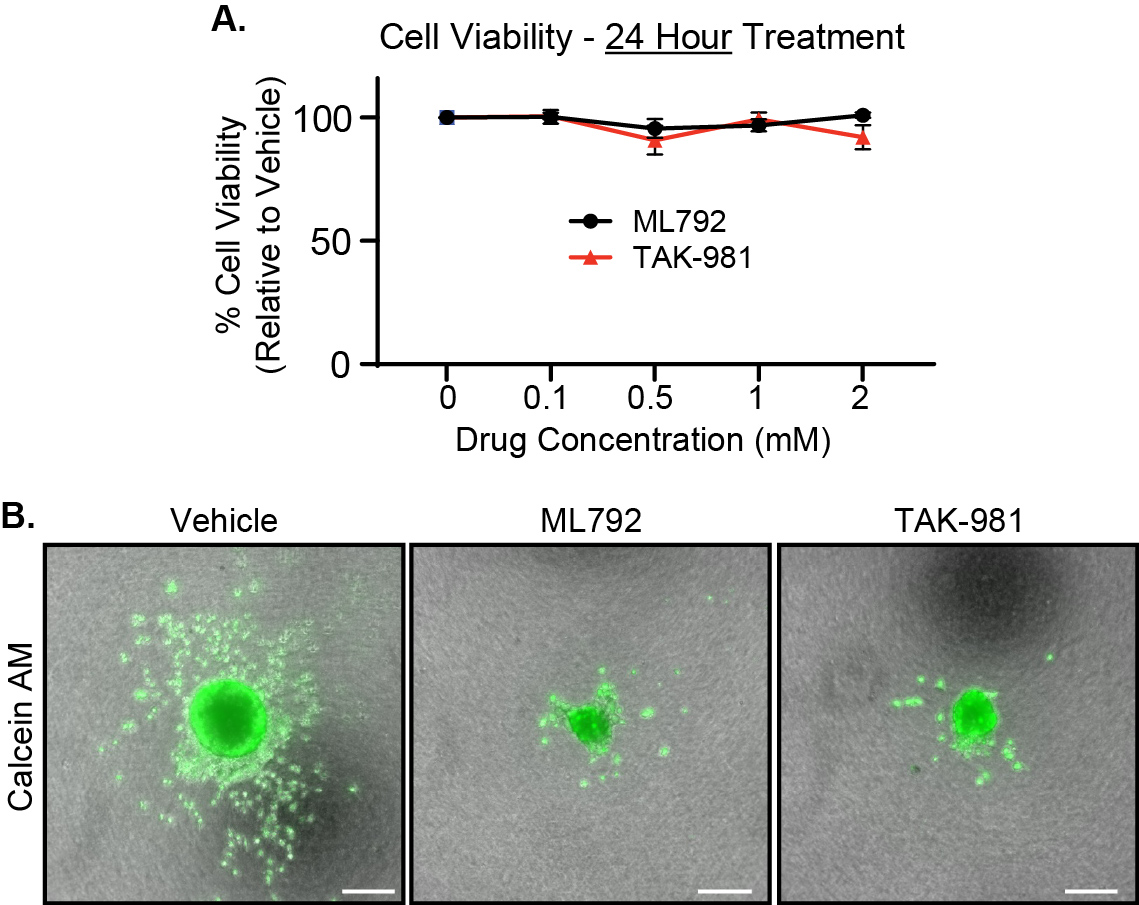

### Supplementary Figure 7

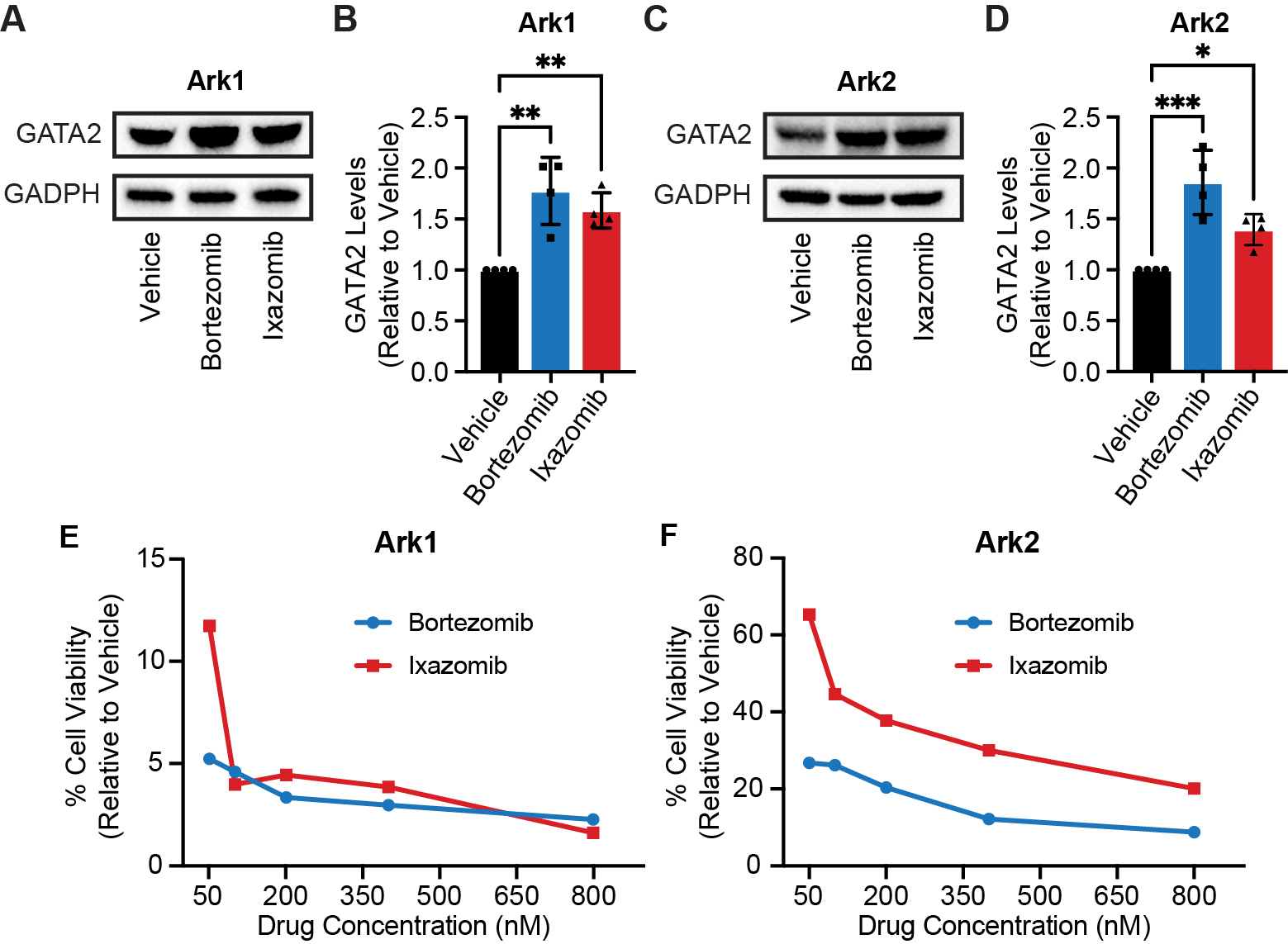

### Supplementary Figure 8

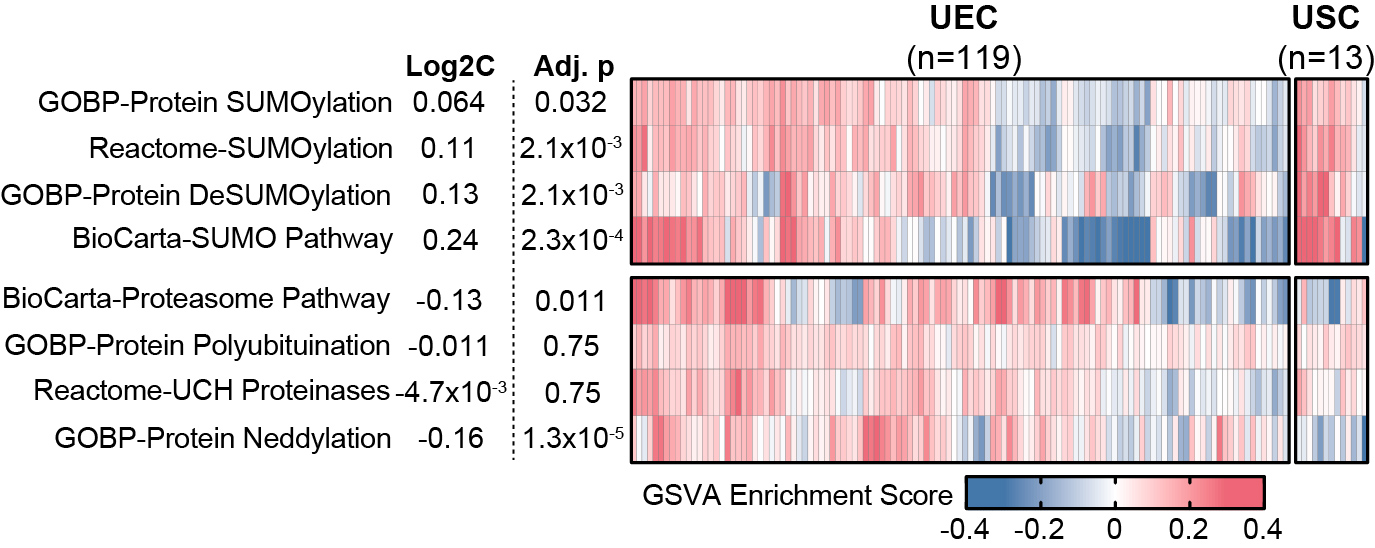
